# The BH3-only protein NOXA serves as an independent predictor of breast cancer patient survival and defines susceptibility to microtubule targeting agents

**DOI:** 10.1101/2021.05.26.445825

**Authors:** Gerlinde Karbon, Manuel D. Haschka, Hubert Hackl, Claudia Soratroi, Lourdes Rocamora-Reverte, Walther Parson, Heidelinde Fiegl, Andreas Villunger

## Abstract

Breast cancer (BC) treatment frequently involves microtubule-targeting agents (MTAs), such as paclitaxel, that arrest cells in mitosis. Sensitivity to MTAs is defined by a subset of pro- and anti-apoptotic BCL2 family proteins controlling mitochondrial apoptosis. Here, we aimed to determine their prognostic value in primary tumour samples from 92 BC patients. Our analysis identified high *NOXA/PMAIP* mRNA expression levels as an independent prognostic marker for improved relapse-free survival (RFS) and overall survival (OS) in multivariate analysis in BC patients, independent of their molecular subtype. Analysis of available TCGA datasets of 1060 BC patients confirmed our results and added a clear predictive value of *NOXA* mRNA levels for patients who received MTA-based therapy. In this TCGA cohort, 122 patients received MTA-treatment and high *NOXA* mRNA levels correlated with their progression-free interval (PFI) and OS. Our follow-up analyses in a panel of BC cell lines of different molecular subtypes identified NOXA protein expression as a key determinant of paclitaxel sensitivity in triple-negative breast cancer (TNBC) cells. Moreover, we noted highest additive effects between paclitaxel and chemical inhibition of BCLX, but not BCL2 or MCL1, documenting dependence of TNBC cells on BCLX for survival and paclitaxel sensitivity, defined by NOXA expression levels.

## INTRODUCTION

BC is with 13.3% the most common type of cancer in women (1). BC can be classified into three main subfamilies according to the presence or absence of the hormone receptors for estrogen (ER) and progesterone (PR) and the human epidermal growth factor receptor 2 (HER2) status: Luminal A/B (about 40%, ER^+^/PR^+/-^/HER2^-^), HER2^+^ (10-15%, ER^-^/PR^-^/HER2^+^) and those negative for all these marker, referred to as TNBC (15-20%, ER^-^/PR^-^/HER2^-^). TNBC therapy involves aggressive chemotherapies due to lack of clear molecular targets (2, 3). Recently PARP1 inhibitors, Olaparib and Talazobarib, and the immune checkpoint inhibitor Atezolizumab, targeting PD-L1, in combination with the microtubule targeting agent (MTA) paclitaxel are used, reviewed by Lyons (4).

MTAs, like vincristine or paclitaxel, inhibit microtubule dynamics (5). This eventually activates the spindle assembly checkpoint (SAC) and triggers mitotic (M)-arrest when applied in tissue culture, eventually leading to apoptosis (6). Paclitaxel shows success in treating metastatic breast and ovarian cancer, as well as various leukaemias (5, 7). Although MTAs are largely successful, resistance and neurotoxicity limit their broader application (8). One way to evade mitotic cell death (MCD) is to overexpress anti-apoptotic BCL2 proteins (9). In BC, MCL1, BCL2 and BCLX are often found amplified (10, 11), making them more resistant to different types of therapeutics (12, 13), including paclitaxel (14). MCD is a desired outcome in cancer therapy, yet clinical efficacy also involves alternative anti-proliferative and pro-inflammatory effects (15). Of note, tumour cells often manage to escape cell death in a process called “mitotic slippage”, the premature exit from mitosis, triggered by the gradual decay of cyclin B levels below a critical threshold, allowing cell survival (16, 17).

We and others recently demonstrated that the molecular mechanism underlying MCD depends on the activity of BH3-only proteins, most notably BIM and NOXA, and the degradation of anti-apoptotic MCL1 (18). We could further demonstrate that NOXA protein mediates the degradation of MCL1 during extended M-arrest and that knockdown of NOXA leads to MCL1 stabilisation and resistance to MTAs in HeLa cervical cancer and A549 lung cancer cells (18). In a follow-up study, we reported that the co-degradation of NOXA/MCL1 complexes during extended M-arrest requires the mitochondrial E3-ligase MARCH5 (19), suggesting that its inhibition may help increase the efficacy of MTAs. Interestingly, ablation of the mitochondrial GTPase DRP1, deregulating mitochondrial network dynamics, sensitizes epithelial cancer cells to MTA-induced apoptosis (20). Taken together, this places mitochondria at the core of mitotic cell death regulation upon MTA treatment success.

Along this line, BH3 mimetics, inhibiting anti-apoptotic BCL2 proteins, are confirmed to be sufficient to prime cancer cells to various chemotherapeutics, including paclitaxel (21). BH3-mimetics bypass the need for upstream inducers of BH3-only proteins, such as p53 or PTEN, which are frequently impaired in human cancers. The first valid prototype of a BH3-mimetic, ABT-737, targeting BCL2, BCLX and BCLW, showed promising efficacy in haematological malignancies (22). However, several cancer types are resistant to ABT-737, or its orally bioavailable successor, Navitoclax (ABT-263), due to the overexpression of MCL1 (23). ABT-199 (Venetoclax), solely targeting BCL2 within the sub-nanomolar-range, shows clinical efficacy in CLL (24), is heavily explored in clinical trials and shows promising results in ER^+^ BC patient-derived xenotransplants (PDX) (25). Wehi-539 and its successors, A-1155463 and A-1331852, are targeting BCLX within sub-nanomolar-range (26) but cause thrombocytopenia, limiting clinical application (27, 28). More recently, a highly specific MCL1 inhibitor, S63845, was shown to have potent anti-tumour activity as a single-agent in preclinical leukaemia models (29), as well as in combination with ABT-199 in BC PDX studies (30).

Here, we tested the predictive value of BCL2 family expression levels for BC patient survival in a patient cohort with detailed clinical follow-up identifying *NOXA* mRNA as an independent prognostic marker. We then expanded our analysis to the TCGA database for confirmation of our findings and investigated the biological significance of the NOXA/MCL1 axis for MCD in BC cell lines exposed to paclitaxel and the additive potential of various BH3-mimetics, highlighting the relevance of BCLX for TNBC cell survival.

## RESULTS

### High *NOXA/PMAIP* mRNA expression predicts superior survival of BC patients receiving MTA-based therapy

Previous analyses of BCL2 family members in BC databases have focused exclusively on the pro-survival BCL2 subset across molecular subtypes (25, 30). We aimed to provide a more comprehensive picture of BCL2 family expression comparing malignant *vs*. non-malignant tissue by including pro-apoptotic members. Therefore we investigated the mRNA expression of pro-apoptotic effectors (BAX, BAK, BOK), BH3-only proteins previously involved in MTA-induced cell death (BID, BIM, PUMA, NOXA) and anti-apoptotic BCL2 family members (MCL1, BCL2, BCLX, BCLW, BCLB) within fresh-frozen tissue samples of different molecular subtypes from 92 patients with primary BC. Our first analysis revealed significant differences in relative mRNA levels between healthy and diseased tissue for anti-apoptotic *BCL2, BCLX/BCL2L1* and *MCL1*. Of note, *BCL2* expression was significantly lower in cancerous tissue (Fig. 1A), while *BCLX* and *MCL1* levels were found increased (Fig. 1B). Analysis of their pro-apoptotic counterparts, the BH3-only proteins *BID* and *NOXA*, revealed a significant increase in mRNA levels in cancerous tissues compared to non-neoplastic tissue (Fig. 1B). In contrast, *PUMA* and *BIM/BCL2L11* levels and the effector proteins *BAX, BAK* or *BOK* or the anti-apoptotic proteins *BCLW* and *BCLB* were comparable (Fig 1C). Pearson correlation analyses revealed significant associations for the co-expression of *MCL1* with *PUMA* or *BAX*, as well as *BCLW/BCL2L2* with *PUMA* or *BAX.* Amongst the pro-death proteins, an interdependence was noted between *BAX* and *BAK* or the BH3-only protein *PUMA* or *BID* with *BAX*. (Supp. Fig. 1), suggesting co-regulation of gene expression.

**Figure 1:**
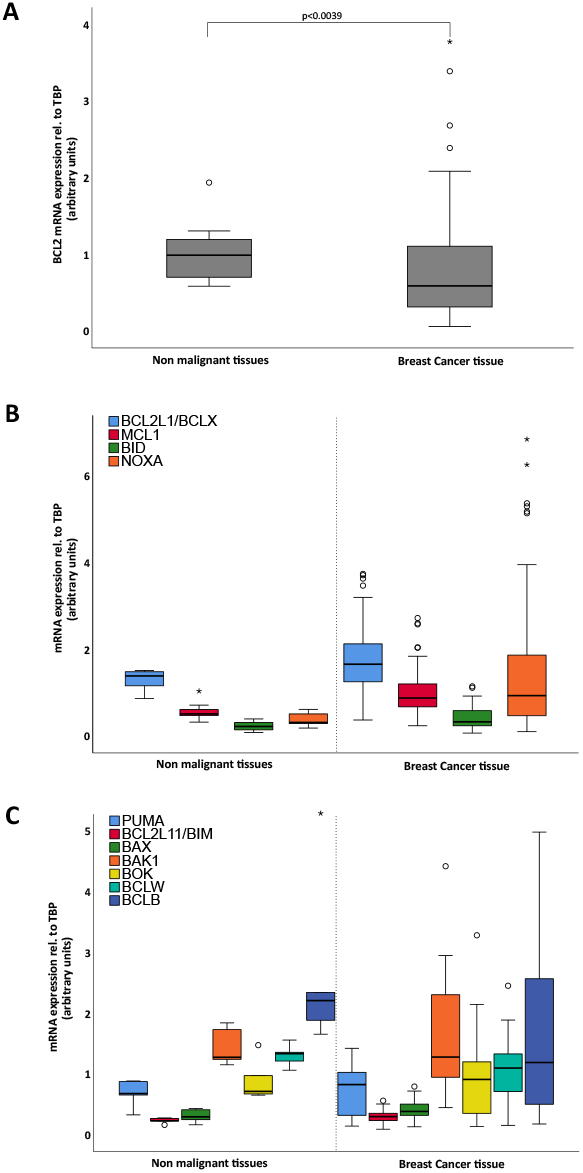
mRNA expression analysis of BCL2 family members in 10 non neoplastic and 92 neoplastic breast tissues. mRNA expression of (**A**) BCL2 (p=0.039), (**B**) BCLX (p=0.035), MCL1 (p=0.001), BID (p=0.020) and NOXA (p=0.002) and (**C**) PUMA, BIM, BAX, BAK, BOK, BCLW and BCLB. Extreme values are marked with asterisks, outliers with circles.

When analyzing clinical parameters, we confirmed previous findings (25) by documenting high *BCL2* mRNA expression in low-grade and luminal A type BC patients still expressing HR (Supp. Table 1). Consistent with BCL2 being a target of HR or signalling, levels were generally higher in HR^+^ tumours (Supp. Table 1). BCLX expression correlated with the same parameters and was enriched in *HER2*^+^ tumours but no longer associated with low tumour grade. Neither *MCL1* nor *BCLW* mRNA showed any correlation with clinical parameters, but *BCLW* showed higher expression in large tumours. Amongst pro-apoptotic genes, expression of *BID* associated with high tumour grade while *NOXA* was found higher expressed in medullary carcinomas (Supp. Table 1).

Performing ROC analyses to define the best cut-off to identify significant patient survival differences across cancer subtypes revealed a clear correlation between higher *BCL2* expression levels and a superior RFS. Higher levels of *MCL1* correlated significantly with a good OS in univariate analyses (Table 1). Of note, *NOXA* and *BOK* expression showed a strong correlation with both RFS and OS. While high *NOXA* expression was associated with superior survival, strong *BOK* expression was associated with poor survival (Table 1). Importantly, this correlation pattern for *BOK* and *NOXA* was maintained when multivariate analyses were performed (Table 2). Expression of *BCL2* still correlated with RFS, while levels of *MCL1* correlated only with improved OS (Table 2). Notably, these correlations were verified by analyzing the TCGA database containing expression data of 1060 BC patients; from those, 471 patients were treated with chemotherapy, from which 112 patients received MTAs (Fig. 2A-D). Strikingly, *NOXA* mRNA expression levels correlated with OS and PFI within BC patients from the TCGA dataset treated with MTA but no other type of chemotherapy (Fig. 2E,F), showing its relevance and confirming the predictive value of our data (Table 2). Within this large validation cohort, BOK expression levels no longer correlated with survival (data not shown).

**Figure 2:**
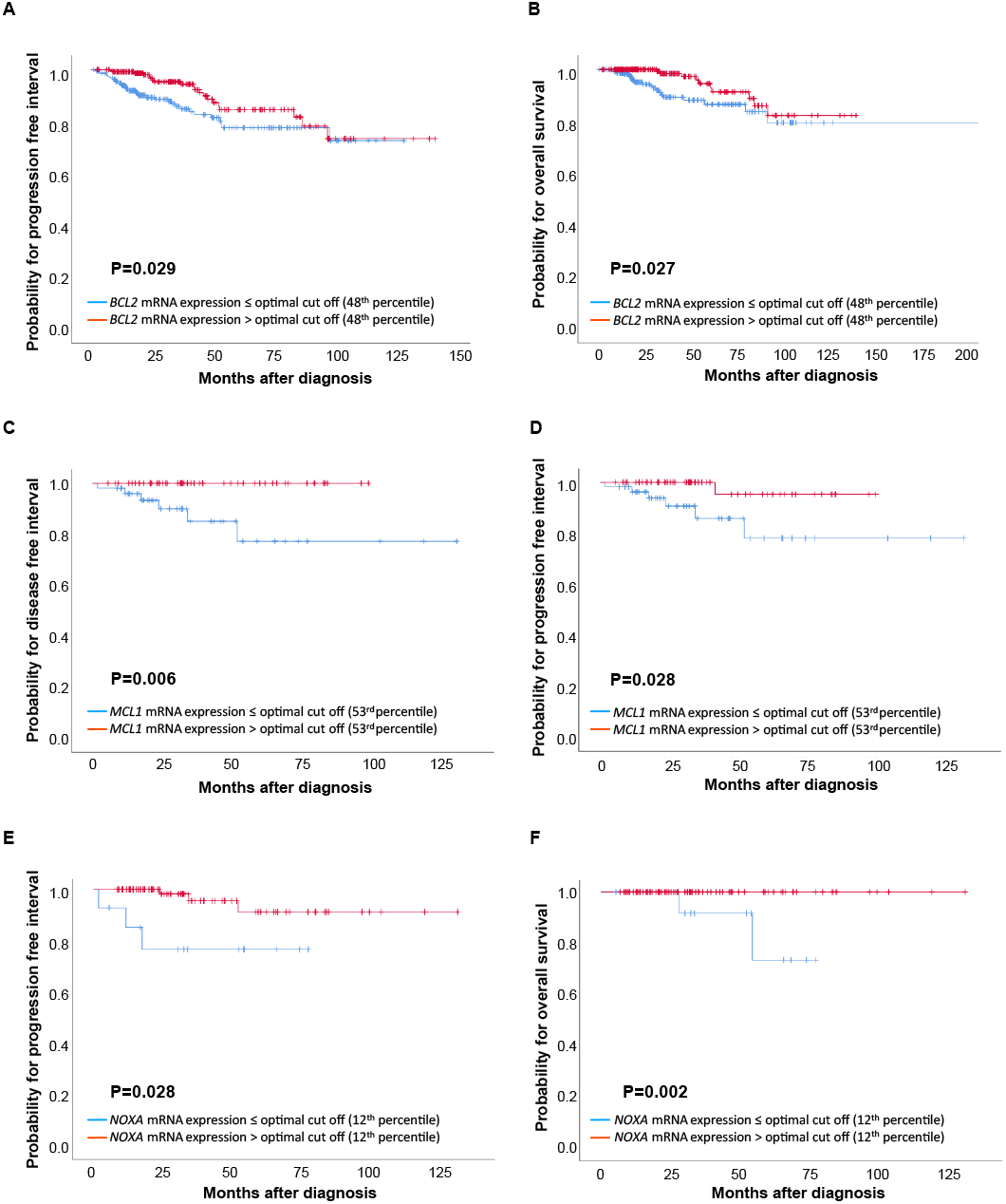
Kaplan Meier survival analysis in the TCGA cohort. (**A**) PFI and (**B**) OS based on BCL2 mRNA-expression in 471 BC patients treated with chemotherapy other than MTA. (**C**) DFI and (**D**) PFI based on MCL1 mRNA-expression in 112 BC patients treated with MTA chemotherapy. (**E**) PFI and (**F**) OS based on NOXA mRNA-expression in 112 BC patients treated with MTA chemotherapy.

**Table 1.** Univariate survival analysis. Relapse free and overall survival in 92 patients with primary breast cancer.

| Variable |  | RELAPSE FREE SURVIVAL |  | OVERALL SURVIVAL |  |
| --- | --- | --- | --- | --- | --- |
|  |  | Median, years (95% CI) | P-value<br>(logrank-Test) | Median, years (95% CI) | P-value<br>(logrank-Test) |
| **Size** | T1 | n.r. | 0.150 | n.r. | 0.093 |
|  | T2/3/4 | 8.09 (0.00 - 19.35) |  | 10.44 (4.62 - 16.25) |  |
| **LN** | negative | n.r. | 0.112 | n.r. | **0.007** |
|  | positive | n.r. |  | 9.93 (6.35 - 13.50) |  |
| **Tumour grade** | I | n.r. | 0.609 | 10.73 (8.69 - 12.78) | 0.886 |
|  | II | 8.09 (n.r.) |  | 14.64 (8.81 - 20.47) |  |
|  | III | n.r. |  | 16.91 (8.76 - 20.06) |  |
| **Histology** | invasive lobular breast cancer | n.r. | 0.187 | 18.69 (5.86 - 31.53) | 0.905 |
|  | invasive ductal breast cancer | 8.09 (0.00 - 17.91) |  | 12.15 (6.51 - 17.79) |  |
|  | medullary breast cancer | n.r. |  | 16.91 (5.20 - 28.62) |  |
| **MP** | premenopausal | 8.09 (0.00 - 19.69) | 0.492 | n.r. | 0.106 |
|  | postmenopausal | n.r. |  | 10.73 (5.33 - 16.14) |  |
| **HER2** | neg | n.r. | 0.127 | 16.91 (12.90 - 20.92) | 0.067 |
|  | pos | 5.23 (3.85 - 6.62) |  | 7.25 (3.22 - 11.28) |  |
| **ER** | neg | n.r. | 0.646 | 16.91 (5.01 - 28.81) | 0.696 |
|  | pos | 19.23 (2.81 - 35.65) |  | 14.64 (8.94 - 20.35) |  |
| **PR** | neg | n.r. | 0.709 | 10.20 (0.00 - 22.45) | 0.376 |
|  | pos | 19.23 (3.15 - 35.32) |  | 14.64 (9.48 - 19.80) |  |
| **MTA chemotherapy** | no | 5.95 (0.00 - 16.78) | 0.057 | 8.76 (6.60 - 10.92) | **0.001** |
|  | yes | n.r. |  | n.r. |  |
| **Radiation therapy** | no | n.r. | 0.316 | 14.64 (9.68 - 19.61) | 0.870 |
|  | yes | 6.77 (0.00 - 17.27) |  | 17.78 (9.74 - 25.83) |  |
| **Endocrine therapy** | no | n.r. | 0.658 | 14.64 (4.38 - 24.90) | 0.601 |
|  | yes | 8.09 (0.00 - 18.43) |  | 12.15 (6.23 - 18.07) |  |
| **BCL2 mRNA expression** | low ($\leq 48^{\text{th}}$ percentile) | 4.72 (2.07 - 7.36) | **0.003** | 8.59 (2.04 - 15.14) | 0.117 |
| | high ($> 48^{\text{th}}$ percentile) | n.r. | | 15.46 (7.41 - 23.52) | |
| **MCL1 mRNA expression** | low ($\leq 53^{\text{rd}}$ percentile) | 5.95 (n.r.) | 0.128 | 9.35 (7.11 - 11.59) | **0.014** |
| | high ($> 53^{\text{rd}}$ percentile) | 19.23 (n.r.) | | 17.95 (n.r.) | |
| **NOXA mRNA expression** | low ($\leq 12^{\text{th}}$ percentile) | 1.98 (1.44 - 2.52) | **<0.001** | 3.28 (1.64 - 4.92) | **<0.001** |
| | high ($> 12^{\text{th}}$ percentile) | n.r. | | 16.91 (13.39 - 20.43) | |
| **BOK mRNA expression** | low ($\leq 34^{\text{th}}$ percentile) | n.r. | **0.008** | n.r. | **0.010** |
| | high ($> 34^{\text{th}}$ percentile) | 5.95 (2.70 - 9.20) | | 10.44 (6.92 - 13.96) | |

*Note:* The significance level (P) was determined by log-rank test.
Abbreviations: LN, lymph node status; MP, menopausal status; HER2, human epidermal growth factor receptor 2 status; ER, oestrogen receptor status; PR, progesterone receptor status; HR, hormone receptor status. n.r. not reached

**Table 2.** Multivariate Cox-Regression survival analysis. Relapse free survival and overall survival in 92 patients with primary breast cancer.

| Variable |  | RELAPSE FREE SURVIVAL |  | OVERALL SURVIVAL |  |
| --- | --- | --- | --- | --- | --- |
|  |  | Relapse-free survival<br>HR of relapse (95% CI) | P-value | Overall survival<br>HR of death (95% CI) | P-value |
| Size | T1 vs. T2/3/4 | 1.63 (0.66 - 4.02) | 0.289 | 1.49 (0.60 - 3.69) | 0.386 |
| LN | neg. vs. pos. | 1.79 (0.75 - 4.29) | 0.193 | 2.91 (1.17 - 7.24) | **0.022** |
| Tumour grade | grade I vs. grade II/III | 1.94 (0.42 - 8.90) | 0.394 | 1.02 (0.34 - 3.10) | 0.975 |
| MP | pre vs. post | 0.63 (0.34 - 1.19) | 0.157 | 1.35 (0.70 - 2.62) | 0.369 |
| HR | neg. vs. pos. | 0.87 (0.42 - 1.80) | 0.709 | 0.80 (0.42 - 1.54) | 0.507 |
| HER2 | neg. vs. pos. | 1.47 (0.74 - 2.92) | 0.269 | 1.76 (0.91 - 3.40) | 0.091 |
| MTA chemotherapy | no vs. yes | 2.00 (0.95 - 4.21) | 0.067 | 3.31 (1.35 - 8.16) | **0.009** |
| **BCL2 mRNA expression** | Low vs. high (<or>48<sup>th</sup> percentile) | 0.31 (0.15 - 0.66) | **0.002** | - | - |
| **MCL1 mRNA expression** | Low vs. high (<or>53<sup>rd</sup> percentile) | - | - | 0.41 (0.22 - 0.79) | **0.008** |
| **NOXA mRNA expression** | Low vs. high (<or>12<sup>th</sup> percentile) | 0.14 (0.06 - 0.32) | **<0.001** | 0.15 (0.06 - 0.38) | **<0.001** |
| **BOK mRNA expression** | Low vs. high (<or>34<sup>th</sup> percentile) | 4.16 (1.74 - 9.93) | **0.001** | 4.47 (2.05 - 9.78) | **<0.001** |

Note : The significance level was determined by Cox regression analysis. HR, hazard ratio.
Abbreviations: LN, lymph node status; MP, menopausal status; HER2, human epidermal growth factor receptor 2 status; RR, relative risk

In summary, *NOXA* expression levels were the most robust predictor of RFS and OS in our cohort and the TCGA database. Moreover, its mRNA expression levels with superior OS, and PFI in the patient group that received MTA-treatment.

### NOXA protein expression defines MTA-sensitivity in TNBC cell lines

We further investigated the expression profile of NOXA and its relevance for MTA-treatment in relation to other members of the BCL2 protein family in more detail. Therefore, we chose eight different BC cell lines, representing the three main subfamilies: Luminal A/B (MCF-7, T47D and ZR-75-1), TNBC (HS-578-T, MDA-MB-231, Cal-51 and BT20) and HER2^+^ (SKBR3) and analysed protein expression levels of the most common pro-and anti-apoptotic proteins.

The expression of BCL2, NOXA, BIM, BCLB and BOK differed substantially amongst cell lines, whereas MCL1, BCLX, or BID expression levels show less variability (Fig. 3A, B). The protein expression of NOXA was highest in the TNBC cell lines, while BOK was hardly detectable in this subset. MDA-MB-231, HS-578-T and T47D also showed a slightly higher expression of BCLX than the other BC cell lines (Fig. 3A), suggesting co-dependence on BCLX for survival. The strong expression of different BCL2 pro-survival proteins suggests variable dependency for cell survival that does not correlate with a particular molecular subtype, which we investigated in the next step using selective BH3-mimetics.

**Figure 3:**
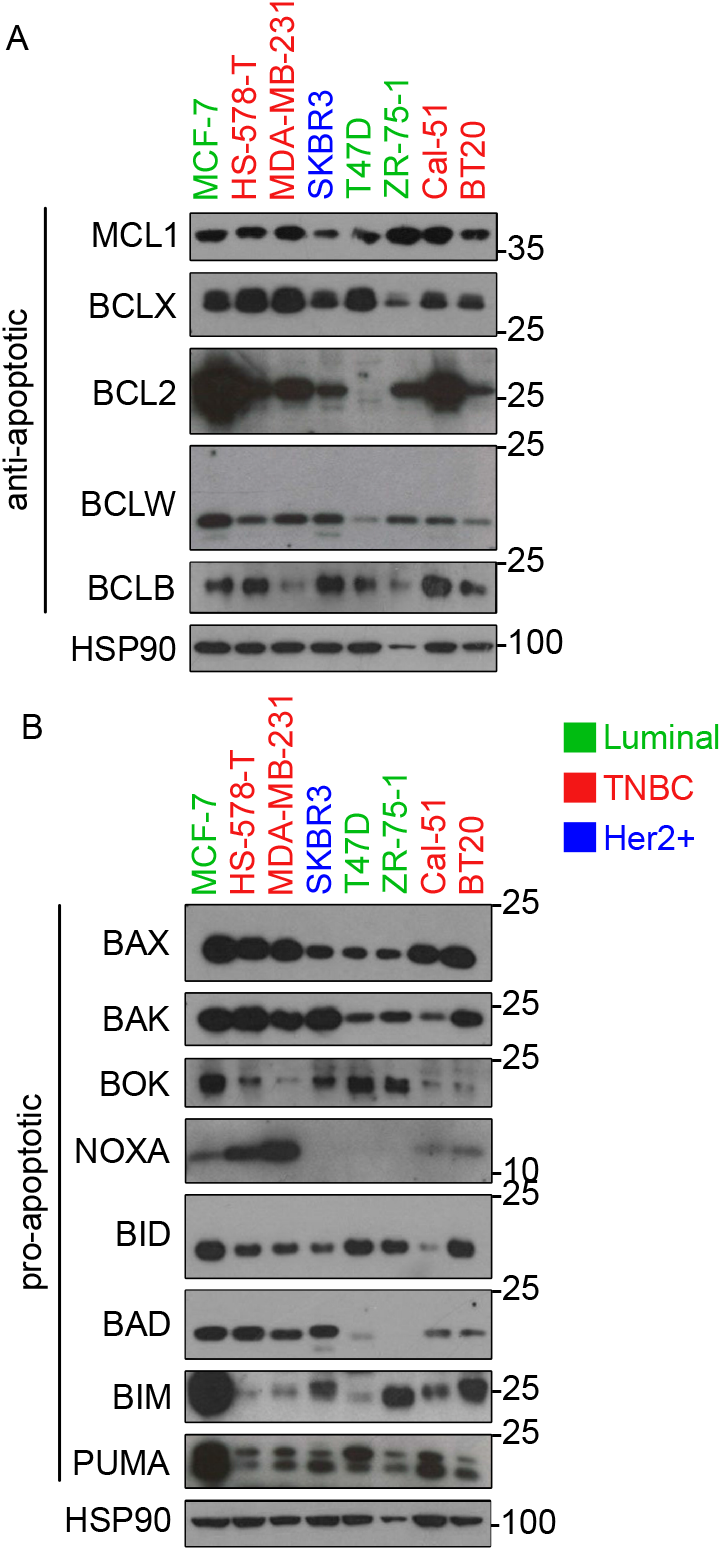
BCL2 protein family expression in different BC cell lines. Different BC cell lines were subjected to western blotting using the indicated antibodies recognizing (**A**) anti- and (**B**) pro-apoptotic BCL-2 proteins. HSP90 was used as a loading control.

### BH3-mimetics efficiently enhance the effects of paclitaxel

All cell lines were treated with graded concentrations of paclitaxel alone or in combination with a fixed concentration of different BH3-mimetics (ABT-737, ABT-199, S63845 and Wehi-539). MTT-assay was used as an indirect readout for cell viability (Supp. Fig. 3). As expected, the cell lines showed different sensitivity against paclitaxel which did not correlate with a particular molecular subtype. Two of the TNBC cell lines, Cal-51 and BT20, were most sensitive to MTA-treatment. Their metabolic activity dropped to about 31% and 25%, respectively, when treated with paclitaxel (Fig. 4A). In contrast, the ER^+^ cell lines ZR-75-1 (83%), T47D (58%) and MCF-7 (50%) showed reduced MTA-sensitivity (Fig. 4A). A saturation using 50 nM paclitaxel was visible for all cell lines tested. Higher concentrations did not reduce metabolic activity any further, with the notable exception of the HS-578-T cells (Supp. Fig. 3).

**Figure 4:**
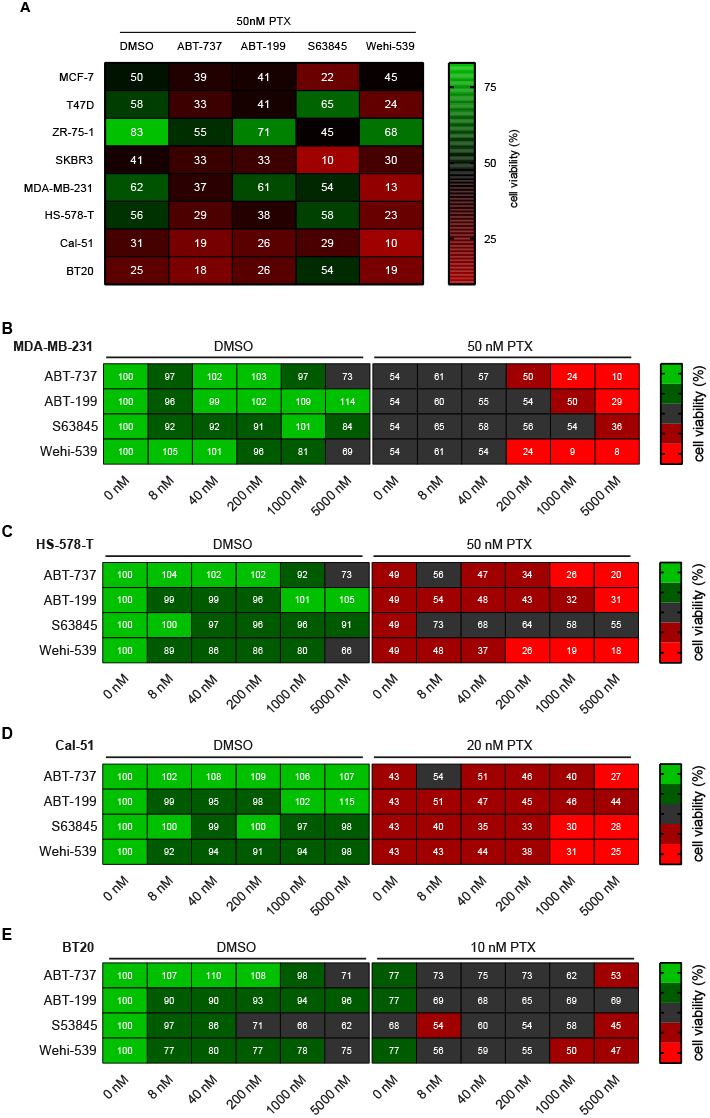
Paclitaxel increases the sensitivity towards BH3-mimetics. (**A**) The indicated cell lines were treated with 50 nM paclitaxel alone or in combination with 1 μM of BH3-mimetics for 48 h. (**B-E**) TNBC cell lines were treated with graded concentrations of BH3-mimetics in combination with a predetermined fixed concentration of paclitaxel for 48h. Metabolic activity was calculated by setting the metabolic activity in relation to the DMSO only control. Heatmaps show mean values of metabolic activity ranging from 0 to 100% as assessed by MTT-assay. (**A**) all cell lines (n=3-4), (**B**) MDA-MB-231 (n=6), (**C**) HS-578-T (n=4), (**D**) Cal-51 (n=6), (**E**) BT20 (n=6).

Inhibiting the pro-survival BCL2 proteins with the different BH3-mimetics alone was mostly ineffective. The BCL2 inhibitor ABT-199 did not affect any cell line at the assayed concentrations (up to 5 μM), including MCF7 cells, which showed the highest BCL2 expression. Similar, the MCL1 inhibitor S63845 only affected MCF-7 and SKBR3 cell lines (Supp. Fig. 2). However, ABT-737, inhibiting BCL2, BCLX and BCLW, and the BCLX inhibitor, Wehi-539, showed activity in some cell lines when used at high doses as a single agent (Fig. 4B-E). Notably, combining paclitaxel with BH3-mimetics led to additive effects, despite variations in their overall potency (Fig. 4, Supp. Fig. 2). In combination with the MCL1 inhibitor S63845, paclitaxel most potently reduced the metabolic activity of MCF7 cells from 50% (paclitaxel) to 22% (paclitaxel + S63845; Fig.4A). The combination of paclitaxel with the BCL2 inhibitor ABT-199 had only a mild effect on HS-578-T or MCF-7 cells despite their high BCL2 levels. (Fig. 4A, Supp. Fig. 2).

Potent effects on all cell lines tested were consistently seen when paclitaxel was combined with ABT-737, having the most substantial impact on the TNBC cell lines HS-578-T and MDA-MB-231. Their metabolic activity was further reduced on average by 27% and 25%, respectively, in the combination setting (Fig. 4A). This finding points towards a critical role for BCLX for cell survival after paclitaxel treatment. Consistently, the most pronounced effects were seen using paclitaxel together with the selective BCLX inhibitor, Wehi-539. This inhibitor further decreased metabolic activity by 33% and 49% in the HS-578-T and MDA-MB-231, respectively (Fig. 4). An increase of the paclitaxel concentration to 500 nM combined with Wehi-539 eliminated all viable MDA-MB-231 cells (Supp. Fig. 3A).

To optimise the effect between paclitaxel and BH3-mimetics, we chose the estimated LD50 concentration of paclitaxel and titrated the different BH3-mimetics. Only inhibition of BCLX, by using ABT-737 or Wehi-539, showed effects as single agents at the highest drug concentration used, *i.e*., 5 μM. (Fig. 4B-E, Supp. Fig. 2). An additive effect was best seen in the TNBC cell lines, identifying BCLX as their primary survival factor that, potentially with the NOXA/MCL1 axis, defines responsiveness to anti-mitotic drug treatment.

### The NOXA/MCL1 axis controls paclitaxel-induced cell death in TNBC cell lines

We could previously show that NOXA driven degradation of MCL1 during extended M-arrest promotes cell death (18). This may explain why chemical inhibition of MCL1 had little effect on paclitaxel sensitivity, as it is automatically degraded upon paclitaxel treatment. We chose the TNBC cell lines HS-578-T, MDA-MB-231 and Cal-51 to analyse the relevance of the NOXA/MCL1 axis for MCD as they showed the strongest NOXA expression. Of note, while MCL1 levels were comparable in these cell lines, NOXA expression appears graded, with the MDA-MB-231 cell line having the highest, followed by HS-578-T and the Cal-51 showing the lowest NOXA levels (Fig. 3).

All three cell lines were synchronised with a double thymidine arrest and released into paclitaxel-containing media to induce prolonged M-arrest. Mitotic shake off was used to enrich cells in M-arrest. CDK1-mediated phosphorylation of CDC27 in mitosis, a component of the anaphase-promoting complex (APC), validated the synchronisation procedure (Fig. 5). We observed that MCL1 and NOXA are co-degraded during extended M-arrest in all three TNBC cell lines. Overall, the NOXA levels followed MCL1 expression and peaked in G2/M before being degraded together (Fig. 5)

**Figure 5:**
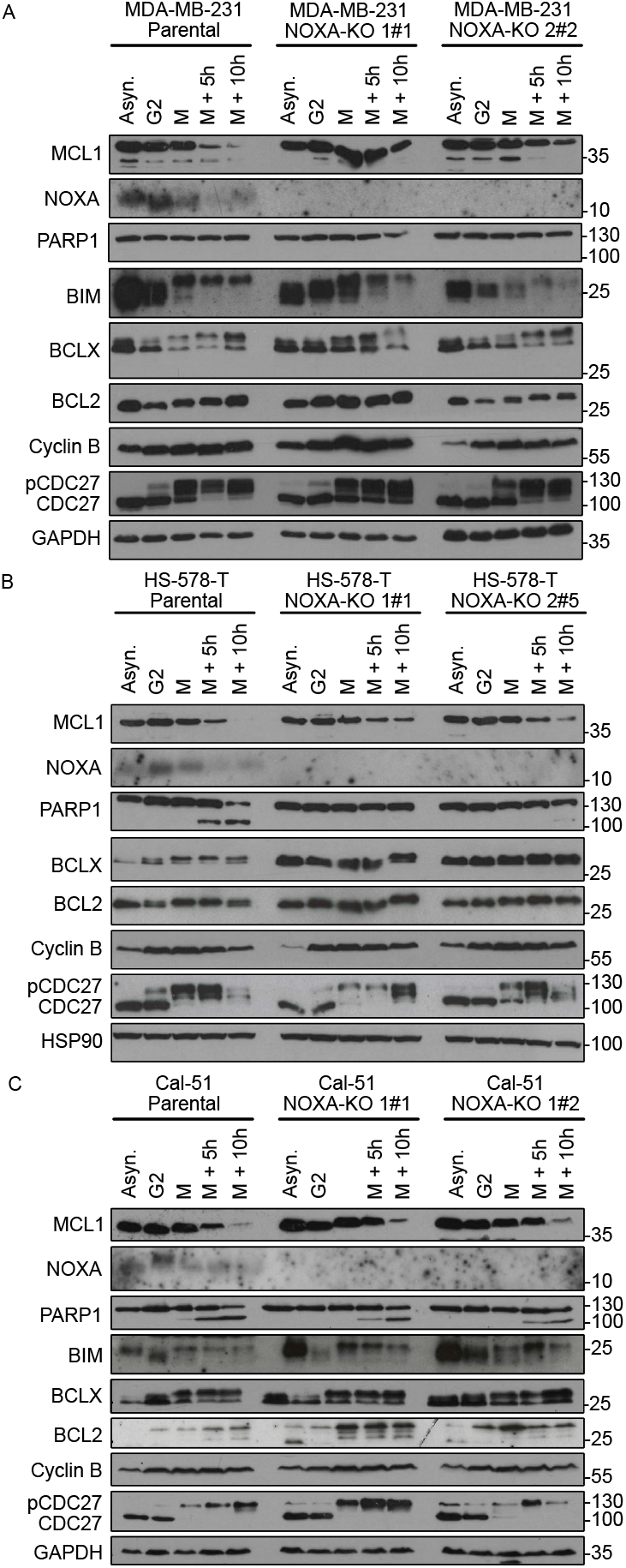
TNBC cell lines treated with PTX show MCL1 and NOXA co-degradation. **(A-C)** TNBC proficient (Parental) or deficient in NOXA (NOXA-KO) were left asynchronous (Asyn.) or synchronized with a double-thymidine block and released into 500 nM PTX. Cells were harvested in G2 and early M-phase (M); part of the M-phase cells was reseeded and harvested 5 h (M +5h) and 10 h (M+ 10h) later. Harvested cells were subjected to western analysis using the indicated antibodies.

The reduction of MCL1/NOXA levels correlated well with apoptosis onset, as monitored by caspase-mediated cleavage of PARP1. Cal-51 cells showed the strongest PARP1 cleavage during extended M-arrest, followed by HS-578-T cells. In strong contrast, MDA-MB-231 cells showed no detectable PARP1 cleavage, indicating resistance against paclitaxel (Fig. 5A). These patterns fit the observed paclitaxel sensitivity/resistance phenotypes noted in the MTT-assays presented above (Fig.4A, Supp Fig.3F).

All three cell lines showed the described phosphorylation of BIM in mitosis, which is needed to promote its APC^CDC20^-driven degradation (18, 31). BCLX and BCL2 are well known to be phosphorylated during mitosis (9, 32-34); this was best observed for BCLX, most of it migrating significantly slower in SDS-PAGE less notable for BCL2, at least with the antibody used (Fig. 5).

To assess the relevance of NOXA/MCL1 turnover for tumour cell survival, we generated NOXA-KO derivatives from these three TNBC cell lines using two independent *sgRNAs* targeting *NOXA*. While the steady-state levels of MCL1 did not substantially differ in asynchronous cells, we could observe a clear stabilisation of MCL1 in the absence of NOXA compared to parental cells upon extended M-arrest (Fig 5). HS-578-T cells showed the most robust MCL1 stabilisation upon M-arrest, followed by the MDA-MB-231 cells, while this effect was modest in the Cal-51 cell line. Regardless, the absence of NOXA was beneficial for survival upon MTA-treatment, as PARP1 cleavage was strongly reduced in the HS-578-T (Fig. 5B) and the Cal-51 cells (Fig. 5C). As there was no PARP1 cleavage detectable in the parental MDA-MB-231 cells, no such effect was observed in NOXA-KO derivative cell lines (Fig 5A).

### NOXA promotes paclitaxel-induced cell death and drives synergy with BH3-mimetics in TNBC cells

Monitoring PARP1 cleavage by western blot may not have been sensitive enough to reveal a contribution of NOXA to paclitaxel-induced cell death, either when used alone or in combination with BH3-mimetics. Hence, to assess if NOXA-deficiency provides MDA-MB-231 cells with a potential survival benefit, we treated these cells with paclitaxel plus BH3-mimetics (Fig. 6). The metabolic activity was reduced in the parental cell line by paclitaxel to a mean of 56%, remaining higher in the NOXA-KO clones (72% and 66%, respectively). This effect became more prominent when paclitaxel was combined with different BH3-mimetics (Fig. 6A, Supp. Fig 4A). Again, ABT-737 and Wehi-539 showed the most potent effects in the parental cell line, indicative of their BCLX dependence in the presence of MTAs. Accordingly, viability was partially rescued by the deletion of NOXA. Similar results were seen in the HS-578-T cell lines upon *NOXA* deletion (Fig. 6B). In the Cal-51 cells, the deletion of *NOXA* conveyed only modest paclitaxel resistance when compared to the other two TNBC cell lines, consistent with the limited impact on PARP1 cleavage noted before (Fig. 5C). This finding goes along with NOXA protein expression being lowest in this cell line and confirms that NOXA function is critical for MCD and acts in good co-operation with BH3-mimetics targeting BCLX.

**Figure 6:**
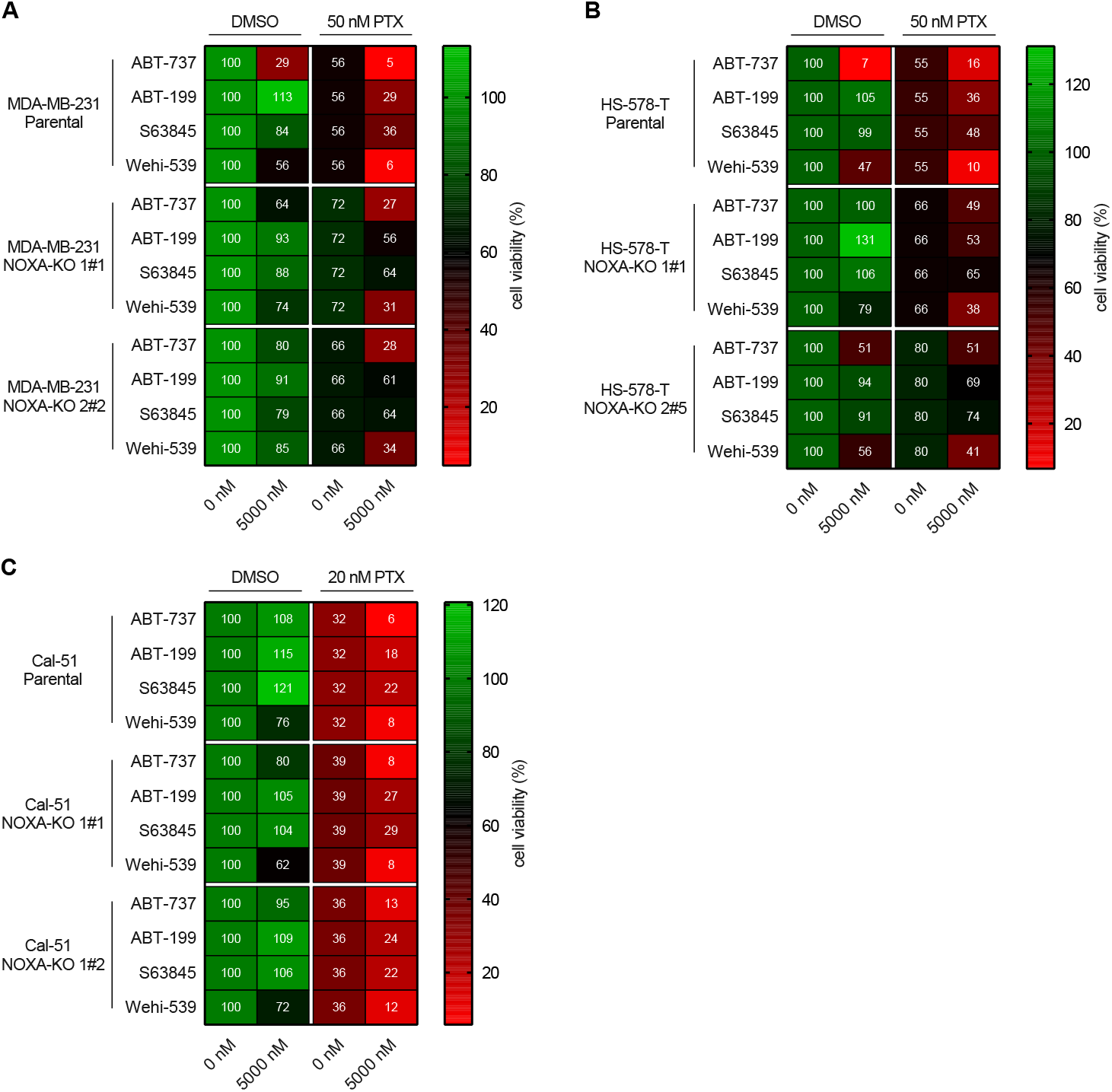
NOXA deletion protects TNBC from PTX and BH3-mimetic co-treatment. Parental and two NOXA-KO clones of each TNBC cell line were treated with or without 5000 nM of the indicated BH3-mimetics alone (DMSO) or in combination with 20 or 50 nM PTX for 48h. Heatmaps are showing the mean value of metabolic activity assessed with MTT-assay. (**A**) MDA-MB-231 (n=5), (**B**) HS-578-T (n=4), (**C**) Cal-51 (n=5).

## DISCUSSION

Analyzing a well-characterized set of 92 BC patient specimens that have subsequently been treated with chemotherapy post-surgery, *NOXA* was identified as the only BH3-only protein with prognostic value across all molecular BC subtypes. High mRNA expression strongly correlated with improved RFS and OS within uni- and multivariate Cox-Regression analysis (Tables 1, 2). This observation could be confirmed in an independent BC patient cohort of the TCGA dataset. Most notably, while our patient cohort size was not large enough to extract statistically robust data on drug-responses, *NOXA* levels also predicted superior PFI and OS in the 122 TCGA patients treated with MTAs, but not for the 350 patients receiving other chemotherapeutic agents (Fig. 2E-F). No such correlation was found for any other BH3-only protein analysed (data not shown).

Of note, mRNA levels of BOK, implicated in endoplasmic reticulum stress induced apoptosis (35), also showed a highly significant prognostic value in uni- and multivariate analysis (Table 1, 2), but not the TCGA data set. BOK has been implicated in Ca^++^ signalling from the ER (36) and pyrimidine synthesis thereby affecting drug resistance and cell proliferation rates (37). Yet, it remains unclear how low BOK expression would benefit BC patients. A more detailed analysis of this protein in BC appears warranted, in particular as loss of BOK reportedly prevents liver cancer in mice (38).

Looking at the predictive value of anti-apoptotic BCL2 proteins, we confirmed high mRNA levels of *BCL2* as beneficial for RFS, as noted before (39, 40), and reconfirmed this within the TCGA dataset (Fig. 2A-B). A similar beneficial effect could be linked to *MCL1* mRNA expression (Fig. 2 C-D). It remains a matter of debate why high levels of a pro-survival protein may improve BC patients’ prognosis. Still, one can imagine a scenario where high BCL2 or MCL1 expression may reduce the pressure to delete other cell death regulators, such as p53, which comes at the price of impaired genomic stability, fostering more aggressive disease (41). In fact, BCL2 overexpression has been shown to delay tumour onset in animal models of irradiation-driven lymphoma and myelodysplastic syndrome transition into AML (42, 43). MCL1 is a short-lived protein regulated mainly at the translational and post-translation level; hence, analysis of protein levels is critical. Consistently, studying a panel of tumour tissue specimens on a tissue microarray revealed that high expression of MCL1 predicts poor outcome in BC in all but the HER2 amplified subtype (44). Moreover, MDA-MB-468 TNBC were highly susceptible to chemical MCL1 inhibition or genetic ablation and tumours forming in the MMTV-PyMT mouse model of BC showed clear MCL1 dependence (44). Consistent with these observations, a recent study reported the beneficial effects of chemical MCL1 inhibition and the MTA docetaxel in TNBC patient-derived xenotransplant (PDX) studies in mice (30). None of these studies addressed the relevance of NOXA for the cell death phenotypes noted.

Our characterisation of the BCL2 protein family expression in cell lines revealed high variation across subtypes (Fig. 3). The low expression of NOXA in SKBR3, T47D and ZR-57-1 (Fig. 3B) might indicate its downregulation as part of a selected survival mechanism. Resistance to therapy is frequently correlated with the downregulation of *NOXA* mRNA in multiple cancer types (45) and linked to the fact that NOXA plays a decisive role in the degradation of MCL1 (18, 46).

The TNBC MDA-MB-231 cell line showed an above-average resistance to paclitaxel, compared to the two other TNBC cell lines. This is consistent with other studies showing high tolerance of MDA-MB-231 against paclitaxel, which may be related to an increased propensity to undergo mitotic slippage (47, 48). Low BAK levels were shown to increase resistance against paclitaxel (49), consistent with MCL1 preferentially binding to BAK (46). We could observe this for the ZR-75-1 and T47D cell line (Fig. 3B, Supp. Fig. 3F, G). However, Cal-51 cells also showed a low BAK expression but are among the most paclitaxel-sensitive cell lines (Fig. 3B, Supp. Fig 3C), suggesting that BAK expression levels alone cannot be seen as a reliable predictor of paclitaxel sensitivity.

Similarly, the expression of BCL2 proteins among our BC cell lines did not allow predictions on their sensitivity against BH3-mimetics, best seen in MCF-7 cells when comparing BCL2 expression with sensitivity to ABT-199. Still, those cells were susceptible to inhibition of MCL1. This indicates that, despite frequent BCL2 upregulation, BCL2 is not a major survival protein for these tumour cells. However, an earlier study using ER^+^ PDX models could show that treating luminal BC with ABT-199 was as effective as ABT-737 combined with chemotherapy (25). This suggests that BCL2 can become a critical survival factor when its co-expressed pro-survival partners are neutralized.

In a genome-wide siRNA screen, MCL1 was shown to be a critical survival factor in TNBC cells (50). We used one of the latest MCL1 inhibitors in clinical development, S63845 (29). However, it showed limited potency against TNBC cell lines when used as a single agent. Yet, the HER2^+^ SKBR3 were highly sensitive to S63845 alone, which correlates with the finding that these cells rely on MCL1 for survival (51). While combining MCL1 inhibition with paclitaxel may be beneficial in some settings (30), our data show that BCLX seems to be generally more critical than MCL1 for BC survival. Application of Wehi-539 or ABT-737 that both target BCLX showed some effect when used as monotherapy, being even more effective when combined with paclitaxel (Fig. 4, Supp. Fig 2), in line with earlier observations (52). BH3-profiling in MDA-MB-231 cells revealed its dependency on BCLX to antagonise pro-apoptotic function (53). However, as a single agent, Wehi-539 was shown to have only a minor effect on the metabolic activity of TNBC, including the MDA-MB-231 (54), fitting our data. Nonetheless, combining Wehi-539 with MTAs in TNBC cell lines, where MCL1 is gradually degraded in a NOXA dependent manner, creates a BCLX dependence (Fig. 5), suggests a potential higher therapy efficacy by the combination of those drugs (55). Consistently, elevated levels of BCL2 and MCL1 can lead to resistance against Wehi-539 (26).

Here, we could show that especially the BCLX inhibitor Wehi-539 sensitized cells to paclitaxel. This phenomenon can be explained in part by the recently noted increase in NOXA levels triggered by MTA-induced micronuclei formation and cGAS-dependent IFN-response, leading to enhanced degradation of MCL1 and BCLX dependence of BC (55). An improved version of Wehi-539, with oral activity, might allow the use of lower doses to avoid thrombocytopenia while still maintaining its anti-cancer efficacy (28). *In vivo* studies of the BCLX inhibitor, A-1331852, already showed an enhancement of the effectiveness of paclitaxel (56), and new PROTAC-based concepts may facilitate clinical application of BCLX inhibition without inducing thrombocytopenia (57, 58).

We could show that deleting NOXA leads to stabilisation of MCL1, giving TNBC a survival benefit during extended M-arrest (Fig. 5), in line with our previous studies in HeLa and A549 cells (18). Stabilised MCL1 can bind BAK and/or sequester BIM, which otherwise would be free to activate the intrinsic apoptotic pathway (46, 59). Upon NOXA dependent degradation of MCL1, more BIM is released from MCL1 sequestration and can execute apoptosis; this can be best seen in Cal-51 NOXA-KO cells (Fig. 5C), which show less protection against M-arrest when compared to MDA-MB-231 NOXA-KO cells. This might rely on the low NOXA expression *per se*, indicating that the NOXA/MCL1 axis might not be that prominent in this cell line or that these cells escape M-arrest by slippage (60). The HS-578-T cell line showed clear survival benefit upon NOXA deletion, as evident by the absent PARP1 cleavage in the NOXA-KO cells. The MDA-MB-231 cell line showed no PARP1 cleavage *per se*, but NOXA deletion still had a positive effect on cell survival, as they were more resistant against paclitaxel and BH3-mimetics co-treatment in long-term assays (Fig. 6).

In summary, the NOXA/MCL1 axis plays a crucial role in TNBC treated with MTAs. Therefore, it could be helpful to increase or restore NOXA expression by, for example, inhibiting its degradation. This would allow a dosage reduction of both MTAs and BCLX inhibitors, thereby avoiding their respective side effects of neurotoxicity and thrombocytopenia (57, 58).

## Material and Methods

### Study design, patients and specimens

Expression levels were quantified by reverse-transcription PCR in mRNA isolated from freshly frozen tumour and adjacent tissue from 92 patients with primary BC treated with chemotherapy after surgery (aged 30.3 to 86.7; median age at diagnosis, 53.0 years) and 10 patients with benign breast diseases (aged 19.8 to 46.0; median age at diagnosis, 35.5 years) treated at the Department of Obstetrics and Gynaecology, Medical University of Innsbruck, Austria. The study was reviewed and approved by the Ethics committee of the Medical University of Innsbruck (reference number: 1021/2017), and was conducted following the Declaration of Helsinki, and in concordance with the Reporting Recommendations for Tumour Marker Prognostic Studies of the National Cancer Institute (REMARK) (61). HR status was identified by immunohistochemistry (IHC).

We analyzed the BC dataset from The Cancer Genome Atlas (TCGA) project (n=1060) described in (62, 63) including OS, DSS data and gene expression data from 471 resected primary breast tumours from patients treated with chemotherapy (aged 26.0 to 84.0 years; median age at diagnosis, 53.0 years).

### RNA isolation and reverse transcription for qRT-PCR

Total cellular RNA extraction, reverse transcription and PCR reactions were performed as previously described (64). Primers and probe for the TATA box-binding protein (TBP; endogenous RNA-control) were used according to Bieche et al. (65). Primerlist in Supp Material Table 3.

### Tissue culture and generation of NOXA KO lines

Cells were cultured in a humidified atmosphere containing 5% CO_2_ at 37°C with the required media (Supp. Material Table 1) and routinely checked for mycoplasma. Amplification of 15 STR and amelogenin loci was carried out in the Institute of Legal Medicine, Innsbruck Medical University (66) for fingerprinting the cell lines in use. Synchronisation with double-thymidine-arrest and generation of NOXA-KO (with CRISPR/Cas9 system) cells was performed as described previously (19); guide sequence in Supp. Material Table 3.

### Metabolic activity assessment

Metabolic activity was determined using the 3-(4,5-dimethylthiazol-2-yl)-2,5-diphenyltetrazolium bromide (MTT)-assay (Cell Proliferation Kit I, Roche Germany, Mannheim) according to manufacturer’s protocol. Cells were seeded in duplicates in 96-well flat-bottom plates and treated 24h later with paclitaxel and/or graded doses of BH3-mimetics. DMSO was used as solvent control. Absorbance was measured at 570 nm and 650 nm using an ELISA plate reader (Sunrise, TECAN) with Magellan software (TECAN, V6.4). Metabolic activity was calculated in Excel, setting the OD-read of DMSO only treated cells to 100% metabolic activity.

### Cell lysis and immunoblot analysis

Cells lysis and immunoblot was performed as previously described (19), used antibodies listed in Supp. Material Table 2.

### Statistical analysis

Mann-Whitney U test was applied to compare mRNA expression levels between groups. Univariate Kaplan-Meier analyses and multivariate Cox survival analyses were used to explore the association of *BCL2* family members mRNA expression levels with survival. Optimal thresholds for survival analyses were identified using Youden’s index (67) based on a receiver operating characteristic (ROC) curve analysis; analyses were performed using SPSS statistical software (version 26.0; SPSS Inc.).

One-tailed paired Student *t*-test was calculated on Prism8 (GraphPad Software) for Supp. Fig. (3). Data in Supp. Fig. (3) are represented with standard errors of the mean (SEM). Statistical significance is shown with symbols: * p-value <0.05, ** p-value <0.01 and *** p-value 0.001.

## Author contribution

GK performed experiments, analyzed data, prepared figures, wrote the manuscript; MH, LRR & CS performed experiments, analyzed data; HH performed bioinformatics analyses; WP performed analysis, HF analyzed patient samples, prepared figures, wrote the manuscript, AV analyzed data, wrote the manuscript, conceived the study.

## Acknowledgements

We are grateful to I. Gaggl and J. Heppke, I. Gaugg and M. Fleischer for excellent technical assistance and all lab members for fruitful discussion. This work was supported by the FWF-funded projects # I-3271 & P29499.

## Abbreviations

BC: Breast cancer
TNBC: Triple-negative breast cancer
HR: Hormone Receptor
ER: Estrogen Receptor
PR: Progesterone Receptor
HER2: Human epidermal growth factor receptor 2
OS: Overall Survival
RFS: Relapse free survival
DFI: disease-free interval
PFI: progression-free interval
M-arrest: Mitotic arrest
MCD: Mitotic cell death
MTA: Microtubule targeting agent
PTX: paclitaxel

## Supplementary Figure Legends

**Supplementary Figure 1:**
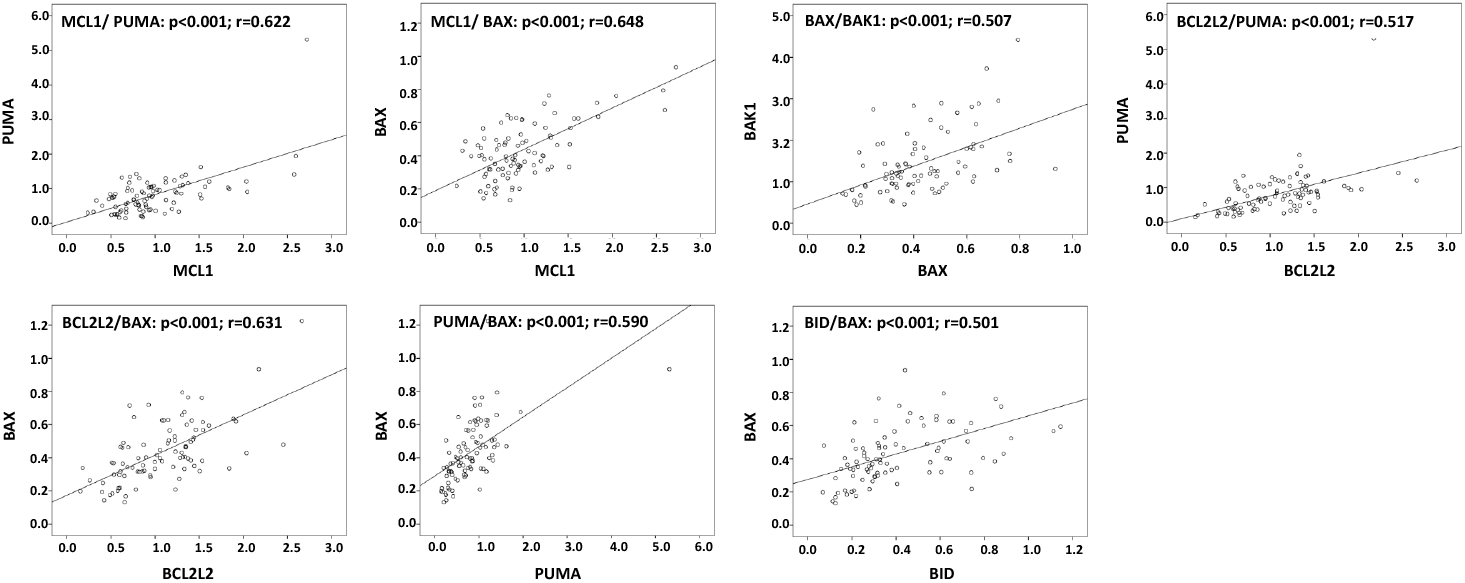
Pearson correlation analysis of BCL2 family member mRNA expression in 92 BC tissues.

**Supplementary Figure 2:**
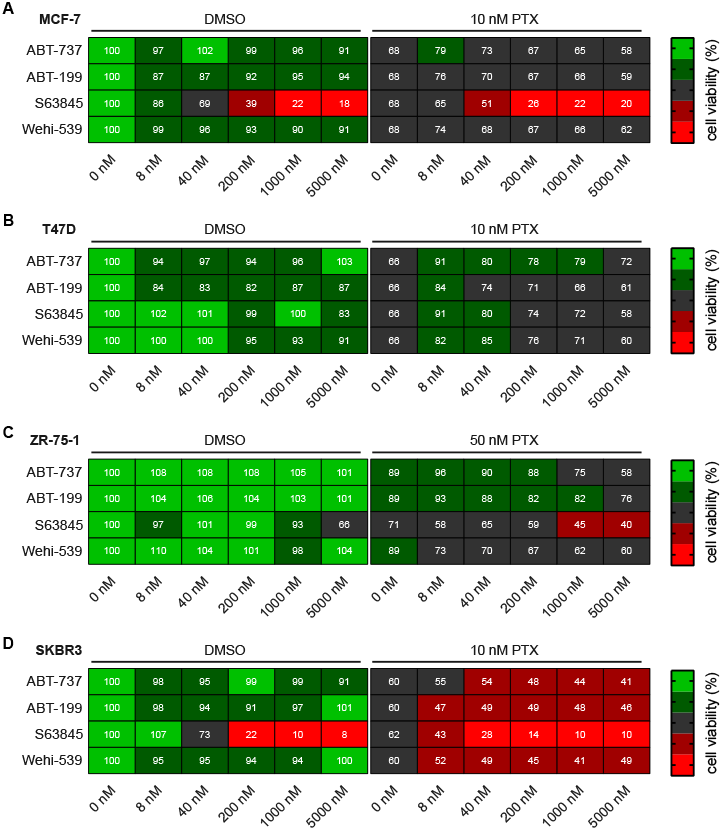
Assessment of metabolic activity with MTT assay after treatment with BH3-mimetics and paclitaxel. (**A-D**) The indicated cell lines were treated with 10 or 50 nM paclitaxel alone or in combination with graded concentrations of different BH3-mimetics (ABT-737, ABT-199, S63845 and Wehi-539) for 48h. DMSO was used as a control. Heatmap is showing the mean value of metabolic activity ranging from 0 to 100% assessed by MTT-assay. (**A**) MCF-7 (n=4), (**B**) T47D (n=4), (**C**) ZR-75-1 (n=4), (**D**) SKBR3 (n=4).

**Supplementary Figure 3:**
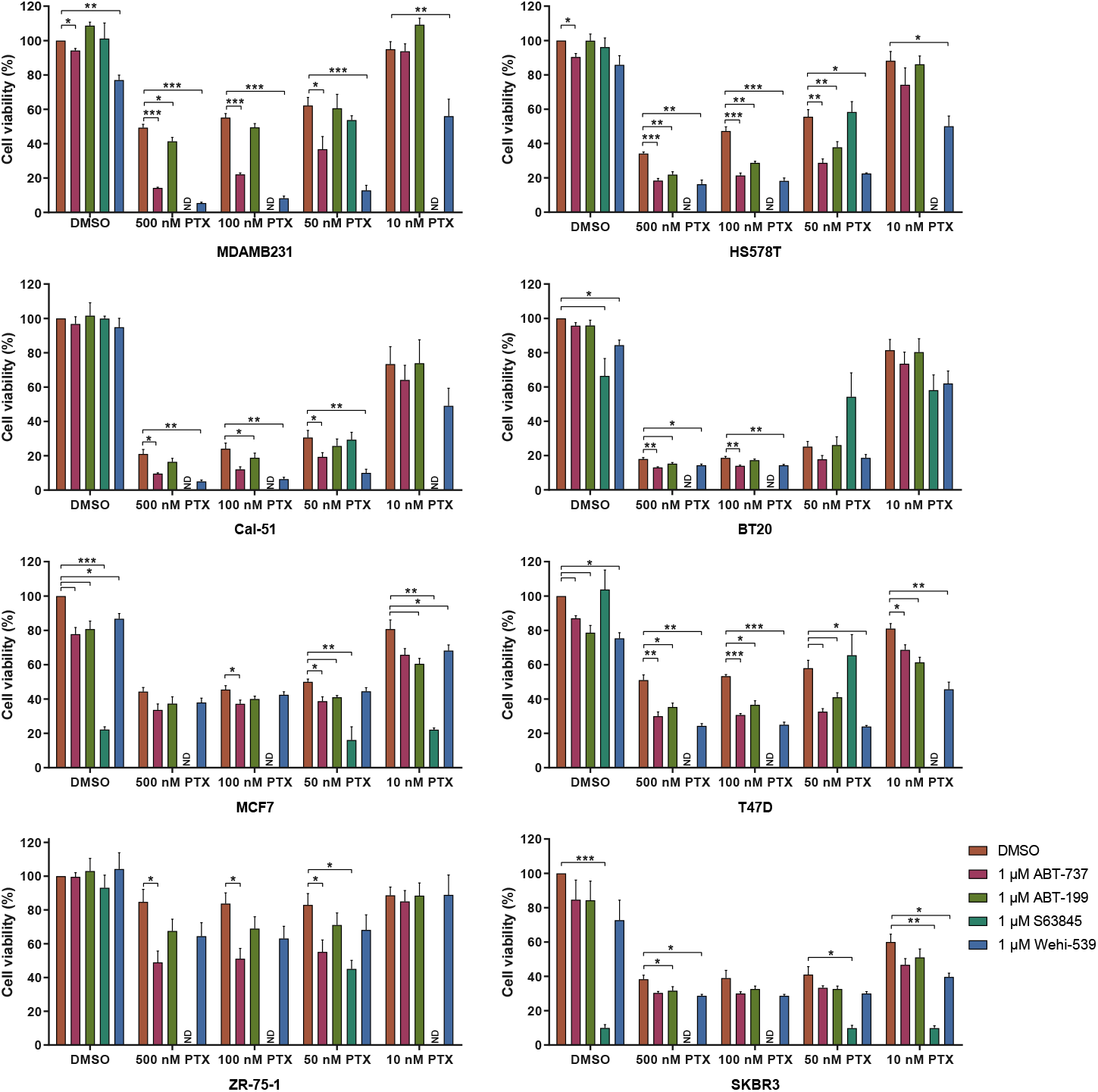
Paclitaxel increases Sensitivity towards BH3-mimetics. (**A-H**) The indicated cell lines were treated with graded doses of paclitaxel in combination with 1 μM of BH3-mimetics for 48 h. Data are shown as mean ±SEM. *: p-value <0.05, **: p-value <0.01,***: p-value <0.001. (**A**) MDA-MB-231(n=4), (**B**) HS-578-T (n=4), (**C**) Cal-51 (n=4), (**D**) BT-20 (n=5), (**E**) MCF-7 (n=4), (**F**) T47D (n=3), (**G**) ZR-75-1 (n=4), (**H**) SKBR3 (n=3).

**Supplementary Figure 4:**
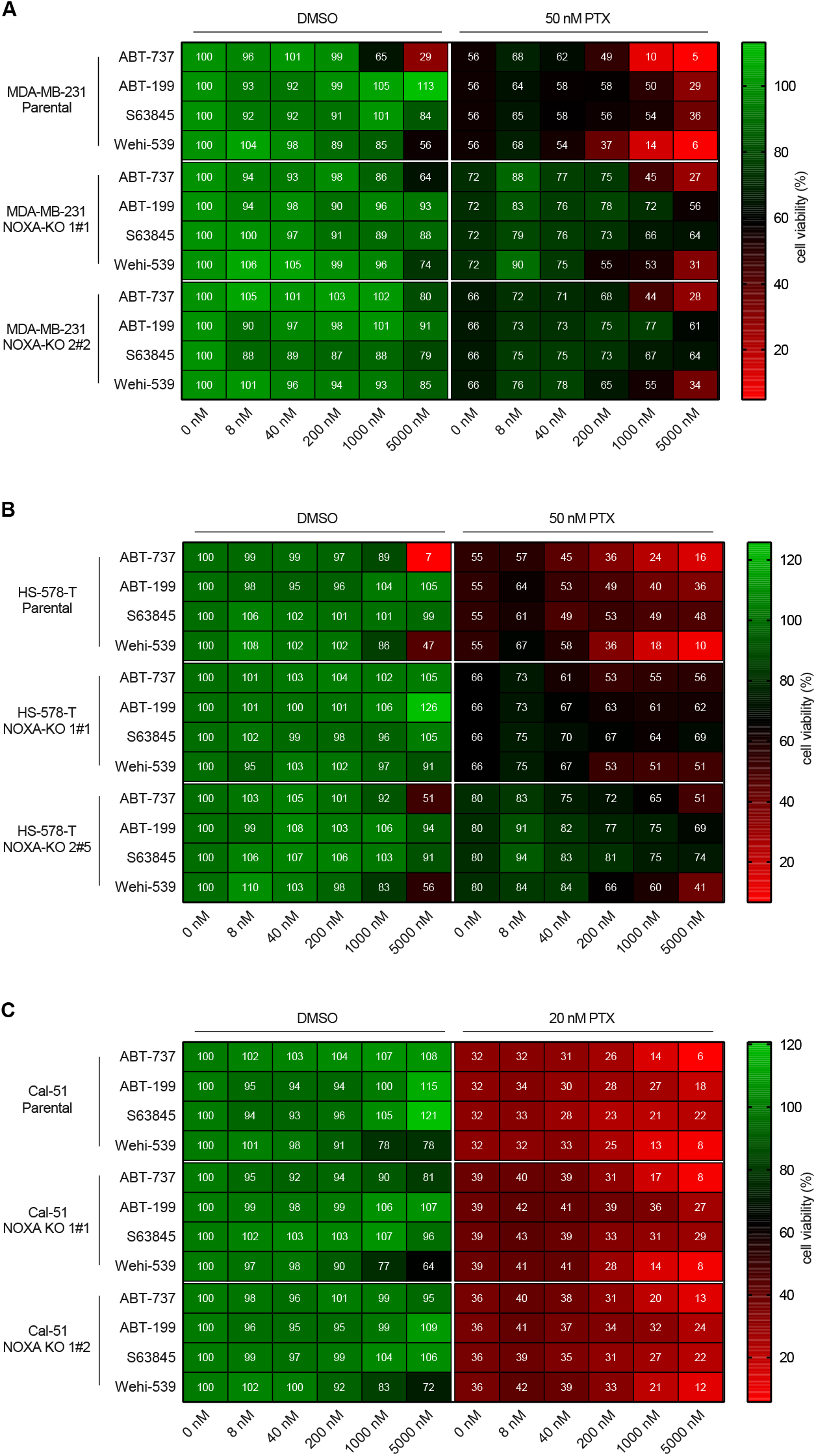
NOXA deletion protects from PTX and BH3-mimetic induced apoptosis. Parental and two NOXA-KO clones of each TNBC cell line were treated with increasing doses of indicated BH3-mimetics alone (DMSO) or in combination with 20 or 50 nM PTX for 48h. Heatmap is showing the mean value of metabolic activity assessed with MTT-assay. (**A**) MDA-MB-231 (n=5), (**B**) HS-578-T (n=4), (**C**) Cal-51 (n=5).

**Supplementary Figure 5:**
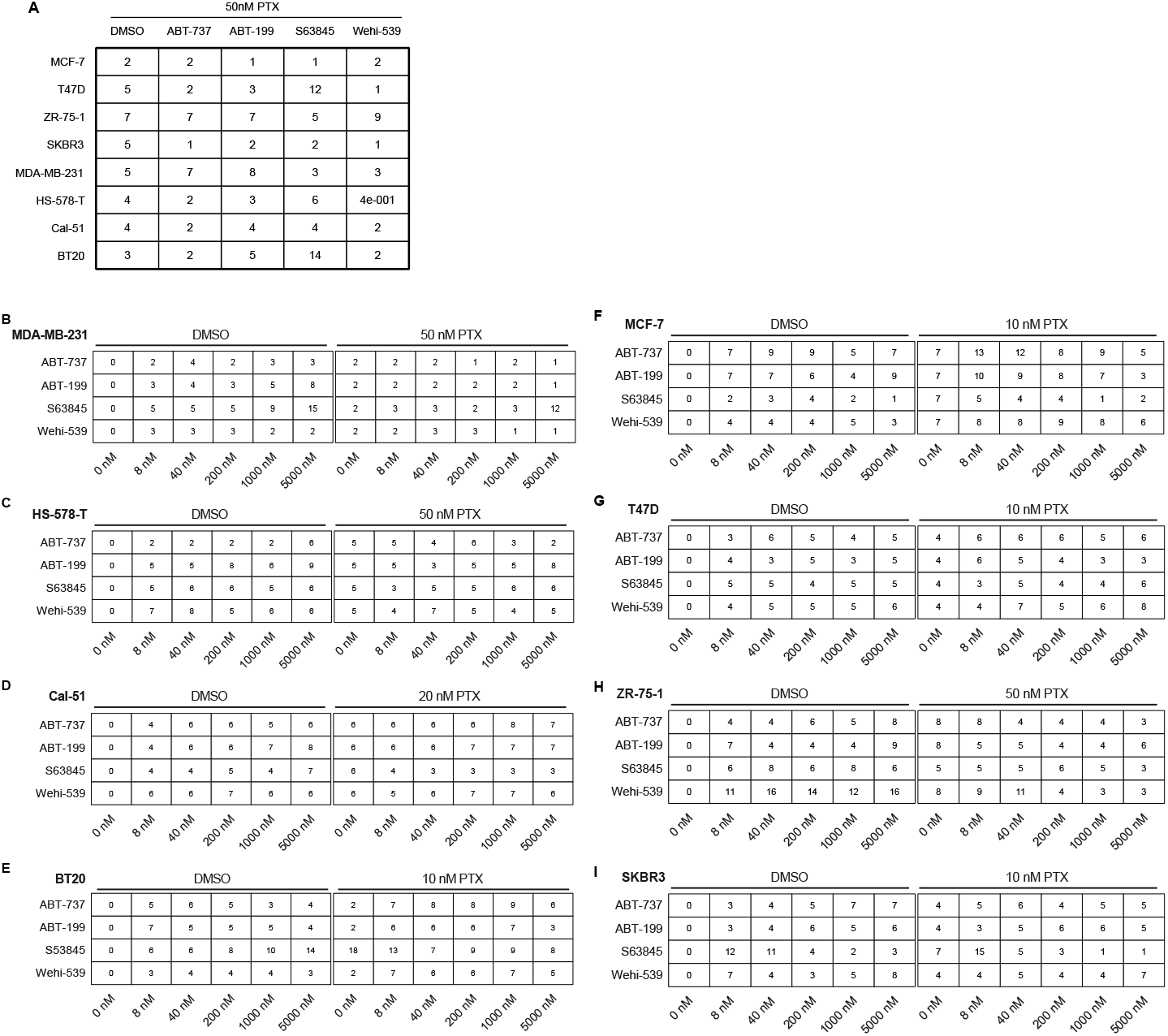
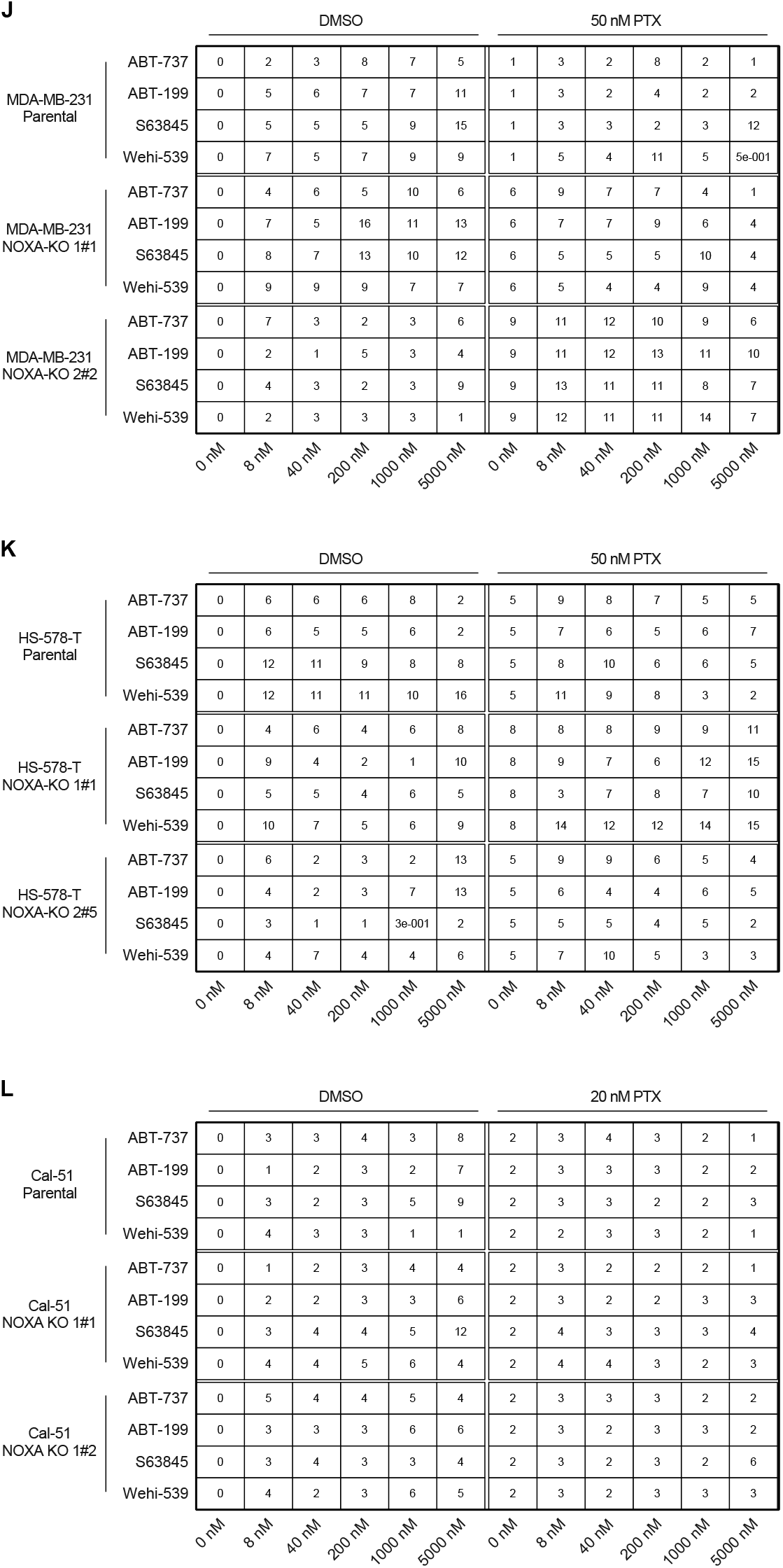
SEM for the heatmaps shown in Fig. 4 (A-E), Supp. Fig. 2 (F-I) and Fig 6 (J-L).

**Supplementary Table 1:**
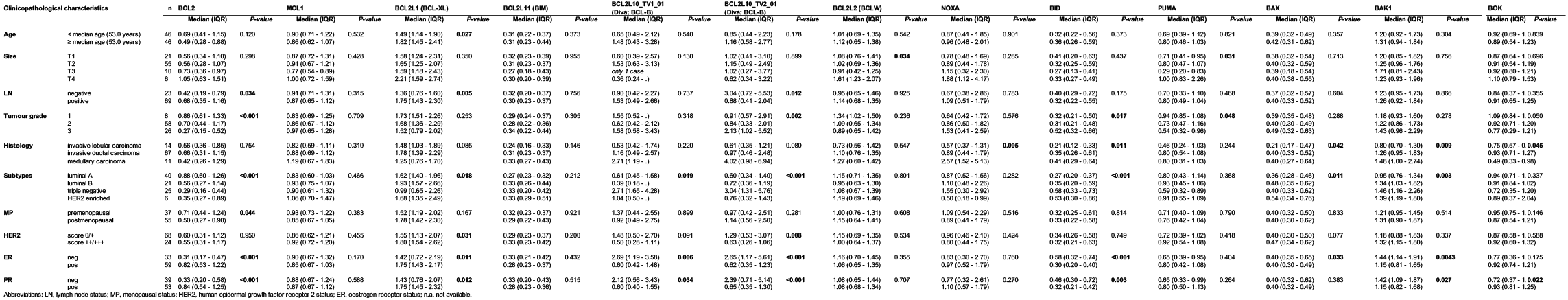
Association of mRNA-expressionof NOXA-MCL1 module associated genes with clinicopathological characteristics in 92 breast cancer patients.

## Supplementary Material

**Table 1.** Media composition for BC cell lines.

**Supplementary Material****Table 1 | Media composition for BC cell lines.**
| Cell line | Media |
| --- | --- |
| **MCF-7** | MEM +Earle (Lonza, BE12-125F)<br>+10% FCS (FBS, PAA laboratories, A15-151)<br>+2 mM L- Glutamin (Sigma G7513)<br>+100 U/ml penicillin and 100 µg/ml streptomycin (Lonza 17-602E)<br>+Non-Essential AA 1x (Gibco 11140-035)<br>+0.01 mg/ml bovine Insulin (sigma-Aldrich I1882) |
| **SKBR3** | Mc Coy's 5A Medium (Sigma Aldrich BE12-688F)<br>+10% FCS (FBS, PAA laboratories, A15-151)<br>+2 mM L- Glutamin (Sigma G7513)<br>+100 U/ml penicillin and 100 µg/ml streptomycin (Lonza 17-602E) |
| **T47D** | RPMI 1640 (Sigma Aldrich, R0883)<br>+10% FCS (FBS, PAA laboratories, A15-151)<br>+2 mM L- Glutamin (PAA laboratories, M11-004)<br>+100 U/ml penicillin and 100 µg/ml streptomycin (Lonza 17-602E) |
| **BT20/**<br>**MDA-MB-231/**<br>**HS-578-T** | EMEM + Earle (Lonza, BE12-125F)<br>+10% FCS (FBS, PAA laboratories, A15-151)<br>+2 mM L- Glutamin (Sigma G7513)<br>+100 U/ml penicillin and 100 µg/ml streptomycin (Lonza 17-602E)<br>+Non-Essential AA 1x (Gibco 11140-035) |
| **ZR-75-1** | RPMI 1640 (Sigma Aldrich, R0883)<br>+10% FCS (FBS, PAA laboratories, A15-151)<br>+2 mM L- Glutamin (Sigma G7513)<br>+100 U/ml penicillin and 100 µg/ml streptomycin (Lonza 17-602E)<br>+4,5 g/l Glucose (Sigma-Aldrich D83752) |
| **Cal-51** | DMEM (Sigma Aldrich, D5671)<br>+10% FCS (FBS, PAA laboratories, A15-151)<br>+2 mM L-Glutamin (Sigma G7513)<br>+100 U/ml penicillin and 100 µg/ml streptomycin (Lonza 17-602E) |

**Table 2.** List of Primary antibodies for Western blot.

| Primary Antibodies |  | Source (dilution) |
| --- | --- | --- |
| Rabbit | MCL1 | Santa Cruz 819 (1:1000) |
| Mouse | NOXA | Calbiochem, clone 114C307 (1:400) |
| Rat | BCLW | Clone 13F9 (Wehi/Alexis) (1:500) |
| Mouse | BCL2 | Clone S100, gift from Andreas Srasser (1 µg/ml) |
| Rabbit | BCLX | Cell Singanlling 2764 (1:1000) |
| Rabbit | BIM | Enzo Life Sciences ADI-AAP-330-E (1:500) |
| Rabbit | PUMA | CS 4976 (1:500) |
| Rabbit | BAD | CS 9292 (1:500) |
| Rabbit | BAX | CS 2772 (1:1000) |
| Rabbit | BAK | CS 3814 (1:1000) |
| Rabbit | BCLB | CS 3869 (1:500) |
| Rat | BID | Clone 8C3 (1:1000) |
| Rabbit | BOK | Clone 1-5-4, gift from Thomas Kaufmann, Bern (1:500) |
| Rabbit | PARP1 | Cell Signalling 9542 (1:1000) |
| Mouse | CDC27 | Clone Bd610455 (1:500) |
| Mouse | Cyclin B | Clone V152 (1:1000) |
| Rabbit | GAPDH | Cell Signalling 2118, clone 14C10 (1:5000) |
| Mouse | HSP90 | Santa Cruz 13119, clone F8 (1:1000) |

**Table 3.**
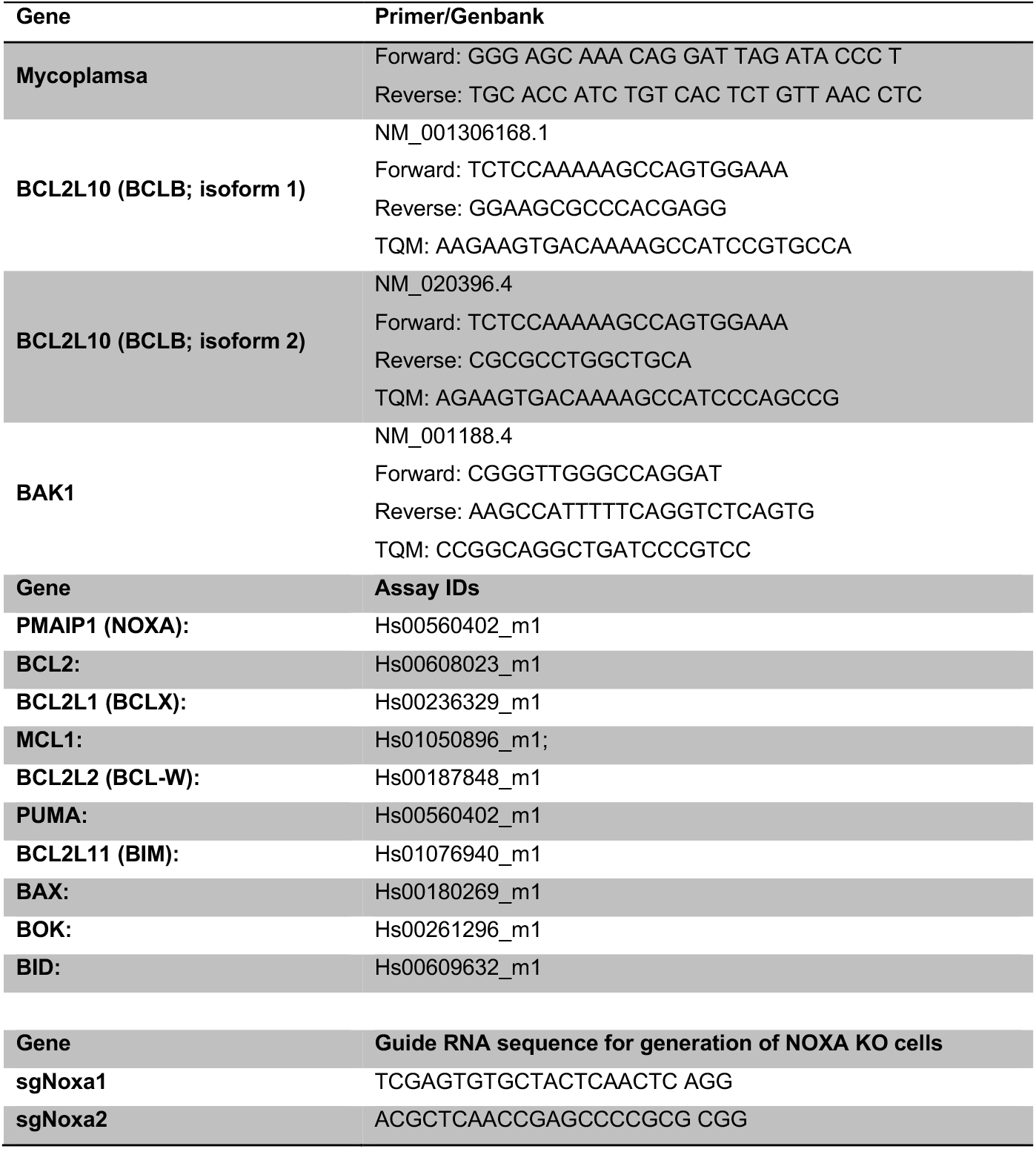
List of Primers. Primers and Probes for BCLB and BAK were determined with the Primer Express software 2.0 (Applied Biosystems, Thermo Scientific, CA, USA). Primers and probe for *BCL2* family members were purchased from Thermo Fisher Scientific (Waltham, Massachusetts, MA, USA, Thermo Fisher Scientific).

**Table 4.** List of Chemical Compounds.

| Compound | Company |
| --- | --- |
| **ABT-737** | Selleck Biochem S1002 |
| **ABT-199** | Apexbio A8194 |
| **S63845** | Selleck Chemicals, 333 S8383 |
| **Wehi-539.** | Apexbio A3935 |
| **Paclitaxel** | Sigma-Aldrich, T7191 |

## References

1. European Commision. 2020 Cancer incidence and mortality in EU-27 countries [Web Page]. 2020 [updated 22.07.2020. Available from: https://ec.europa.eu/jrc/en/news/2020-cancer-incidence-and-mortality-eu-27-countries.

2. Tan DS, Marchio C, Jones RL, Savage K, Smith IE, Dowsett M, et al. Triple negative breast cancer: molecular profiling and prognostic impact in adjuvant anthracycline-treated patients. Breast cancer research and treatment. 2008;111(1):27–44.

3. Waks AG, Winer EP. Breast Cancer Treatment: A Review. Jama. 2019;321(3):288–300.

4. Lyons TG. Targeted Therapies for Triple-Negative Breast Cancer. Current Treatment Options in Oncology. 2019;20(11):82.

5. Čermák V, Dostál V, Jelínek M, Libusová L, Kovář J, Rösel D, et al. Microtubule-targeting agents and their impact on cancer treatment. European journal of cell biology. 2020;99(4):151075.

6. Rieder CL, Maiato H. Stuck in division or passing through: what happens when cells cannot satisfy the spindle assembly checkpoint. Developmental cell. 2004;7(5):637–51.

7. Dumontet C, Jordan MA. Microtubule-binding agents: a dynamic field of cancer therapeutics. Nature reviews Drug discovery. 2010;9(10):790–803.

8. Frederiks CN, Lam SW, Guchelaar HJ, Boven E. Genetic polymorphisms and paclitaxel-or docetaxel-induced toxicities: A systematic review. Cancer treatment reviews. 2015;41(10):935–50.

9. Eichhorn JM, Sakurikar N, Alford SE, Chu R, Chambers TC. Critical role of anti-apoptotic Bcl-2 protein phosphorylation in mitotic death. Cell death & disease. 2013;4:e834.

10. Beroukhim R, Mermel CH, Porter D, Wei G, Raychaudhuri S, Donovan J, et al. The landscape of somatic copy-number alteration across human cancers. Nature. 2010;463(7283):899–905.

11. Williams MM, Lee L, Hicks DJ, Joly MM, Elion D, Rahman B, et al. Key Survival Factor, Mcl-1, Correlates with Sensitivity to Combined Bcl-2/Bcl-xL Blockade. Molecular cancer research: MCR. 2017;15(3):259–68.

12. Perciavalle RM, Opferman JT. Delving deeper: MCL-1’s contributions to normal and cancer biology. Trends in cell biology. 2013;23(1):22–9.

13. Cory S, Huang DC, Adams JM. The Bcl-2 family: roles in cell survival and oncogenesis. Oncogene. 2003;22(53):8590–607.

14. Real PJ, Sierra A, De Juan A, Segovia JC, Lopez-Vega JM, Fernandez-Luna JL. Resistance to chemotherapy via Stat3-dependent overexpression of Bcl-2 in metastatic breast cancer cells. Oncogene. 2002;21(50):7611–8.

15. Weaver BA. How Taxol/paclitaxel kills cancer cells. Mol Biol Cell. 2014;25(18):2677–81.

16. Haschka M, Karbon G, Fava LL, Villunger A. Perturbing mitosis for anti-cancer therapy: is cell death the only answer? EMBO reports. 2018;19(3).

17. Gascoigne KE, Taylor SS. Cancer cells display profound intra- and interline variation following prolonged exposure to antimitotic drugs. Cancer Cell. 2008;14(2):111–22.

18. Haschka MD, Soratroi C, Kirschnek S, Hacker G, Hilbe R, Geley S, et al. The NOXA-MCL1-BIM axis defines lifespan on extended mitotic arrest. Nature communications. 2015;6:6891.

19. Haschka MD, Karbon G, Soratroi C, O’Neill KL, Luo X, Villunger A. MARCH5-dependent degradation of MCL1/NOXA complexes defines susceptibility to antimitotic drug treatment. Cell death and differentiation. 2020;27(8):2297–312.

20. Peña-Blanco A, Haschka MD, Jenner A, Zuleger T, Proikas-Cezanne T, Villunger A, et al. Drp1 modulates mitochondrial stress responses to mitotic arrest. Cell death and differentiation. 2020;27(9):2620–34.

21. D’Aguanno S, Del Bufalo D. Inhibition of Anti-Apoptotic Bcl-2 Proteins in Preclinical and Clinical Studies: Current Overview in Cancer. Cells. 2020;9(5).

22. Chauhan D, Velankar M, Brahmandam M, Hideshima T, Podar K, Richardson P, et al. A novel Bcl-2/Bcl-X(L)/Bcl-w inhibitor ABT-737 as therapy in multiple myeloma. Oncogene. 2007;26(16):2374–80.

23. Yecies D, Carlson NE, Deng J, Letai A. Acquired resistance to ABT-737 in lymphoma cells that up-regulate MCL-1 and BFL-1. Blood. 2010;115(16):3304–13.

24. Souers AJ, Leverson JD, Boghaert ER, Ackler SL, Catron ND, Chen J, et al. ABT-199, a potent and selective BCL-2 inhibitor, achieves antitumor activity while sparing platelets. Nature medicine. 2013;19(2):202–8.

25. Vaillant F, Merino D, Lee L, Breslin K, Pal B, Ritchie ME, et al. Targeting BCL-2 with the BH3 mimetic ABT-199 in estrogen receptor-positive breast cancer. Cancer Cell. 2013;24(1):120–9.

26. Lessene G, Czabotar PE, Sleebs BE, Zobel K, Lowes KN, Adams JM, et al. Structure-guided design of a selective BCL-X(L) inhibitor. Nat Chem Biol. 2013;9(6):390–7.

27. Tao ZF, Hasvold L, Wang L, Wang X, Petros AM, Park CH, et al. Discovery of a Potent and Selective BCL-XL Inhibitor with in Vivo Activity. ACS medicinal chemistry letters. 2014;5(10):1088–93.

28. Wang L, Doherty GA, Judd AS, Tao ZF, Hansen TM, Frey RR, et al. Discovery of A-1331852, a First-in-Class, Potent, and Orally-Bioavailable BCL-X(L) Inhibitor. ACS medicinal chemistry letters. 2020;11(10):1829–36.

29. Kotschy A, Szlavik Z, Murray J, Davidson J, Maragno AL, Le Toumelin-Braizat G, et al. The MCL1 inhibitor S63845 is tolerable and effective in diverse cancer models. Nature. 2016;538(7626):477–82.

30. Merino D, Whittle JR, Vaillant F, Serrano A, Gong JN, Giner G, et al. Synergistic action of the MCL-1 inhibitor S63845 with current therapies in preclinical models of triple-negative and HER2-amplified breast cancer. Science translational medicine. 2017;9(401).

31. Graos M, Almeida AD, Chatterjee S. Growth-factor-dependent phosphorylation of Bim in mitosis. The Biochemical journal. 2005;388(Pt 1):185–94.

32. Choi HJ, Zhu BT. Role of cyclin B1/Cdc2 in mediating Bcl-XL phosphorylation and apoptotic cell death following nocodazole-induced mitotic arrest. Mol Carcinog. 2014;53(2):125–37.

33. Upreti M, Galitovskaya EN, Chu R, Tackett AJ, Terrano DT, Granell S, et al. Identification of the major phosphorylation site in Bcl-xL induced by microtubule inhibitors and analysis of its functional significance. The Journal of biological chemistry. 2008;283(51):35517–25.

34. Deng X, Gao F, Flagg T, May WS, Jr. Mono- and multisite phosphorylation enhances Bcl2’s antiapoptotic function and inhibition of cell cycle entry functions. Proc Natl Acad Sci U S A. 2004;101(1):153–8.

35. Carpio MA, Michaud M, Zhou W, Fisher JK, Walensky LD, Katz SG. BCL-2 family member BOK promotes apoptosis in response to endoplasmic reticulum stress. Proc Natl Acad Sci U S A. 2015;112(23):7201–6.

36. D’Orsi B, Engel T, Pfeiffer S, Nandi S, Kaufmann T, Henshall DC, et al. Bok Is Not Pro-Apoptotic But Suppresses Poly ADP-Ribose Polymerase-Dependent Cell Death Pathways and Protects against Excitotoxic and Seizure-Induced Neuronal Injury. The Journal of neuroscience: the official journal of the Society for Neuroscience. 2016;36(16):4564–78.

37. Srivastava R, Cao Z, Nedeva C, Naim S, Bachmann D, Rabachini T, et al. BCL-2 family protein BOK is a positive regulator of uridine metabolism in mammals. Proc Natl Acad Sci U S A. 2019;116(31):15469–74.

38. Rabachini T, Fernandez-Marrero Y, Montani M, Loforese G, Sladky V, He Z, et al. BOK promotes chemical-induced hepatocarcinogenesis in mice. Cell death and differentiation. 2018;25(4):708–20.

39. Eom YH, Kim HS, Lee A, Song BJ, Chae BJ. BCL2 as a Subtype-Specific Prognostic Marker for Breast Cancer. Journal of breast cancer. 2016;19(3):252–60.

40. Labi V, Erlacher M. How cell death shapes cancer. Cell death & disease. 2015;6(3):e1675.

41. Gurova KV, Kwek SS, Koman IE, Komarov AP, Kandel E, Nikiforov MA, et al. Apoptosis inhibitor as a suppressor of tumor progression: expression of Bcl-2 eliminates selective advantages for p53-deficient cells in the tumor. Cancer biology & therapy. 2002;1(1):39–44.

42. Slape CI, Saw J, Jowett JB, Aplan PD, Strasser A, Jane SM, et al. Inhibition of apoptosis by BCL2 prevents leukemic transformation of a murine myelodysplastic syndrome. Blood. 2012;120(12):2475–83.

43. Michalak EM, Vandenberg CJ, Delbridge AR, Wu L, Scott CL, Adams JM, et al. Apoptosis-promoted tumorigenesis: gamma-irradiation-induced thymic lymphomagenesis requires Puma-driven leukocyte death. Genes Dev. 2010;24(15):1608–13.

44. Campbell KJ, Dhayade S, Ferrari N, Sims AH, Johnson E, Mason SM, et al. MCL-1 is a prognostic indicator and drug target in breast cancer. Cell death & disease. 2018;9(2):19.

45. Montero J, Gstalder C, Kim DJ, Sadowicz D, Miles W, Manos M, et al. Destabilization of NOXA mRNA as a common resistance mechanism to targeted therapies. Nat Commun. 2019;10(1):5157.

46. Willis SN, Chen L, Dewson G, Wei A, Naik E, Fletcher JI, et al. Proapoptotic Bak is sequestered by Mcl-1 and Bcl-xL, but not Bcl-2, until displaced by BH3-only proteins. Genes Dev. 2005;19(11):1294–305.

47. Tabuchi Y, Matsuoka J, Gunduz M, Imada T, Ono R, Ito M, et al. Resistance to paclitaxel therapy is related with Bcl-2 expression through an estrogen receptor mediated pathway in breast cancer. International journal of oncology. 2009;34(2):313–9.

48. Flores ML, Castilla C, Ávila R, Ruiz-Borrego M, Sáez C, Japón MA. Paclitaxel sensitivity of breast cancer cells requires efficient mitotic arrest and disruption of Bcl-xL/Bak interaction. Breast cancer research and treatment. 2012;133(3):917–28.

49. Miller AV, Hicks MA, Nakajima W, Richardson AC, Windle JJ, Harada H. Paclitaxel-induced apoptosis is BAK-dependent, but BAX and BIM-independent in breast tumor. PloS one. 2013;8(4):e60685.

50. Petrocca F, Altschuler G, Tan SM, Mendillo ML, Yan H, Jerry DJ, et al. A genome-wide siRNA screen identifies proteasome addiction as a vulnerability of basal-like triple-negative breast cancer cells. Cancer Cell. 2013;24(2):182–96.

51. Campone M, Noel B, Couriaud C, Grau M, Guillemin Y, Gautier F, et al. c-Myc dependent expression of pro-apoptotic Bim renders HER2-overexpressing breast cancer cells dependent on anti-apoptotic Mcl-1. Molecular cancer. 2011;10:110.

52. Bah N, Maillet L, Ryan J, Dubreil S, Gautier F, Letai A, et al. Bcl-xL controls a switch between cell death modes during mitotic arrest. Cell death & disease. 2014;5:e1291.

53. Ryan JA, Brunelle JK, Letai A. Heightened mitochondrial priming is the basis for apoptotic hypersensitivity of CD4+ CD8+ thymocytes. Proc Natl Acad Sci U S A. 2010;107(29):12895–900.

54. Goodwin CM, Rossanese OW, Olejniczak ET, Fesik SW. Myeloid cell leukemia-1 is an important apoptotic survival factor in triple-negative breast cancer. Cell death and differentiation. 2015;22(12):2098–106.

55. Lohard S, Bourgeois N, Maillet L, Gautier F, Fétiveau A, Lasla H, et al. STING-dependent paracriny shapes apoptotic priming of breast tumors in response to anti-mitotic treatment. Nat Commun. 2020;11(1):259.

56. Leverson JD, Phillips DC, Mitten MJ, Boghaert ER, Diaz D, Tahir SK, et al. Exploiting selective BCL-2 family inhibitors to dissect cell survival dependencies and define improved strategies for cancer therapy. Science translational medicine. 2015;7(279):279ra40.

57. He Y, Zhang X, Chang J, Kim HN, Zhang P, Wang Y, et al. Using proteolysis-targeting chimera technology to reduce navitoclax platelet toxicity and improve its senolytic activity. Nat Commun. 2020;11(1):1996.

58. Kolb R, De U, Khan S, Luo Y, Kim MC, Yu H, et al. Proteolysis-targeting chimera against BCL-X(L) destroys tumor-infiltrating regulatory T cells. Nat Commun. 2021;12(1):1281.

59. Willis SN, Fletcher JI, Kaufmann T, van Delft MF, Chen L, Czabotar PE, et al. Apoptosis initiated when BH3 ligands engage multiple Bcl-2 homologs, not Bax or Bak. Science. 2007;315(5813):856–9.

60. Topham CH, Taylor SS. Mitosis and apoptosis: how is the balance set? Current opinion in cell biology. 2013;25(6):780–5.

61. McShane LM, Altman DG, Sauerbrei W, Taube SE, Gion M, Clark GM, et al. REporting recommendations for tumour MARKer prognostic studies (REMARK). British journal of cancer. 2005;93(4):387–91.

62. Koboldt DC, Fulton RS, McLellan MD, Schmidt H, Kalicki-Veizer J, McMichael JF, et al. Comprehensive molecular portraits of human breast tumours. Nature. 2012;490(7418):61–70.

63. Liu J, Lichtenberg T, Hoadley KA, Poisson LM, Lazar AJ, Cherniack AD, et al. An Integrated TCGA Pan-Cancer Clinical Data Resource to Drive High-Quality Survival Outcome Analytics. Cell. 2018;173(2):400–16.e11.

64. Müller HM, Fiegl H, Goebel G, Hubalek MM, Widschwendter A, Müller-Holzner E, et al. MeCP2 and MBD2 expression in human neoplastic and non-neoplastic breast tissue and its association with oestrogen receptor status. British journal of cancer. 2003;89(10):1934–9.

65. Bièche I, Franc B, Vidaud D, Vidaud M, Lidereau R. Analyses of MYC, ERBB2, and CCND1 genes in benign and malignant thyroid follicular cell tumors by real-time polymerase chain reaction. Thyroid: official journal of the American Thyroid Association. 2001;11(2):147–52.

66. Parson W, Kirchebner R, Mühlmann R, Renner K, Kofler A, Schmidt S, et al. Cancer cell line identification by short tandem repeat profiling: power and limitations. FASEB journal: official publication of the Federation of American Societies for Experimental Biology. 2005;19(3):434–6.

67. Youden WJ. Index for rating diagnostic tests. Cancer. 1950;3(1):32–5.

